# A genetically encoded bifunctional enzyme mitigates redox imbalance and lipotoxicity via engineered Gro3P-Glycerol shunt

**DOI:** 10.1101/2025.06.02.657195

**Authors:** Xingxiu Pan, Subrata Munan, Austin L. Zuckerman, Andrew Pon, Sara Violante, Justin R. Cross, Hardik Shah, Valentin Cracan

**Affiliations:** Scintillon Institute, Laboratory of Redox Biology and Metabolism, San Diego, CA 92121, USA; Program in Mathematics and Science Education, University of California San Diego and San Diego State University, San Diego, CA 92120, USA; Donald B. and Catherine C. Marron Cancer Metabolism Center, Memorial Sloan Kettering Cancer Center, New York, NY 10021, USA; Metabolomics Platform, University of Chicago Medicine Comprehensive Cancer Center, Chicago, IL 60637, USA; Department of Chemistry, The Scripps Research Institute, La Jolla, CA 92037, USA

## Abstract

Dihydroxyacetone phosphate (DHAP), glycerol-3-phosphate (Gro3P) and reduced/oxidized nicotinamide adenine dinucleotide (NADH/NAD^+^) are key metabolites of the Gro3P shuttle system that forms a redox circuit, allowing transfer of reducing equivalents between cytosol and mitochondria. Targeted activation of Gro3P biosynthesis was recently identified as a promising strategy to alleviate reductive stress by promoting NAD^+^ recycling, including in cells with an impaired mitochondrial complex I. However, because Gro3P constitutes the backbone of triglycerides under some circumstances, its accumulation can lead to excessive fat deposition. Here, we present the development of a novel genetically encoded tool based on a di-domain glycerol-3-phosphate dehydrogenase from algae *Chlamydomonas reinhardtii* (*Cr*GPDH), which is a bifunctional enzyme that can recycle NAD^+^ while converting DHAP to Gro3P. In addition, this enzyme possesses an N-terminal domain which cleaves Gro3P into glycerol and inorganic phosphate (Pi) (in humans and other organisms, this reaction is catalyzed by a separate glycerol-3-phosphate phosphatase, a reaction also known as “glycerol shunt”). When expressed in mammalian cells, *Cr*GPDH diminished Gro3P levels and boosted the TCA cycle and fatty acid β-oxidation in mitochondria. *Cr*GPDH expression alone supported proliferation of HeLa cells under conditions of either inhibited activity of the mitochondrial electron transport chain or hypoxia. Moreover, human kidney cancer cells, which exhibit abnormal lipid accumulation, had decreased triglycerides levels when expressing *Cr*GPDH. Our findings suggest that the coordinated boosting of both Gro3P biosynthesis and glycerol shunt may be a viable strategy to alleviate consequences of redox imbalance and associated impaired lipogenesis in a wide repertoire of conditions, ranging from primary mitochondrial diseases to obesity, type 2 diabetes, and metabolic dysfunction-associated steatotic liver disease (MASLD).

## INTRODUCTION

Maintaining the redox balance of NADH/NAD⁺ coenzymes is fundamental to energy metabolism^1^. Glycolysis and the tricarboxylic acid (TCA) cycle utilize oxidized NAD⁺ as an electron acceptor to break down carbohydrates and generate NADH, while the mitochondrial electron transport chain (ETC) oxidizes NADH back to NAD⁺, coupling this process to ATP synthesis^2^. However, NADH produced by cytosolic glycolysis cannot directly enter mitochondria for further oxidation by the ETC. The malate-aspartate shuttle and glycerol-3-phosphate (Gro3P) shuttle are two key redox circuits that help transfer reducing equivalents from cytosol into mitochondria, enabling efficient energy production by the mitochondrial ETC^1,2^. The Gro3P shuttle operates through interconversion of dihydroxyacetone phosphate (DHAP) and Gro3P (**Figure 1a**). Cytosolic Gro3P dehydrogenase (GPD1/GPD1l or cGPDH) converts DHAP into Gro3P while regenerating NAD^+^. Gro3P is then converted back to DHAP by mitochondrial Gro3P dehydrogenase (GPD2 or mGPDH), which faces the mitochondrial intermembrane space and donates electrons to the mitochondrial coenzyme Q pool (**Figure 1a**)^1,2^. Cytosolic NAD^+^ can also be regenerated via pyruvate fermentation, a process catalyzed by lactate dehydrogenase (LDH) which converts pyruvate into lactate^1,2^. Both processes ensure the regeneration of cytosolic NAD^+^ to sustain glycolysis and maintain energy production^3^. Disruptions in these systems can lead to imbalances, such as NADH-reductive stress (increased NADH/NAD^+^ ratio) and Gro3P accumulation^3,4^.

**Figure 1:**
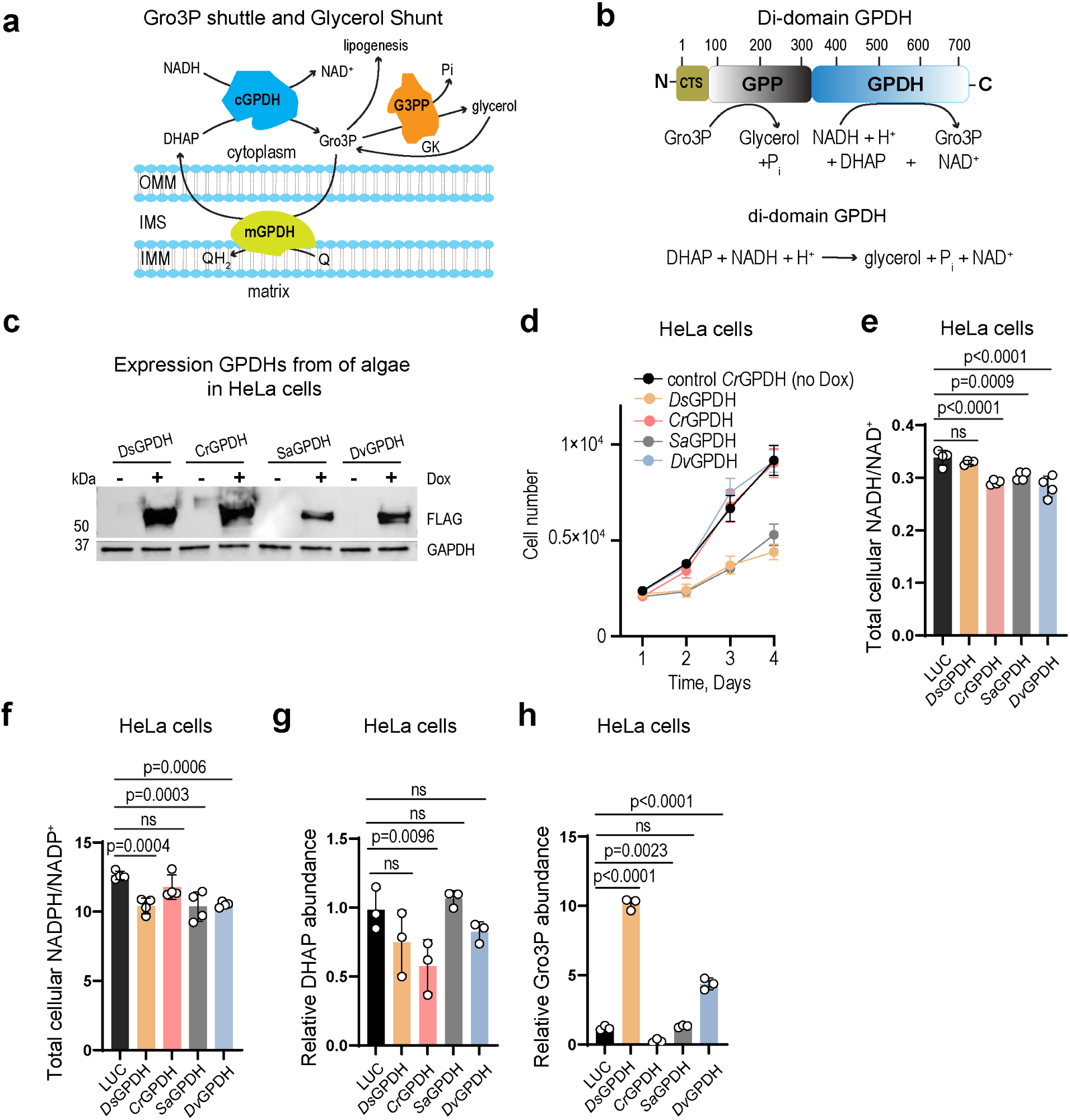
Initial screening of Di-domain GPDHs in mammalian cells. (**a**) Schematic of the mammalian Glycerol-3-phosphate (Gro3P) shuttle and the glycerol shunt. Cytosolic NAD-linked glycerol 3-phosphate dehydrogenase (cGPDH); coenzyme Q-linked mitochondrial glycerol 3-phosphate dehydrogenase (mGPDH); glycerol-3-phosphate phosphatase (G3PP) and glycerol kinase (GK). (**b**) Domain organization and reaction catalyzed by di-domain glycerol-3-phosphate dehydrogenase (GPDH) from algae. CTS, chloroplast targeting sequence; GPP, Gro3P phosphatase domain; GPDH, Gro3P dehydrogenase domain. (**c**) Western blot of *D. salina, C. reinhardtii, S. arctica* and *D. viridis* GPDHs expressed in HeLa cells after 24-hour induction with doxycycline (Dox). A representative western blot is shown. (**d**) The effect of expression of GPDHs from *D. salina, C. reinhardtii, S. arctica* and *D. viridis* on proliferation of HeLa cells grown in pyruvate-free DMEM supplemented with dialyzed FBS (DMEM^+dFBS^). The total cellular NADH/NAD^+^ (**e**) and NADPH/NADP^+^ (**f**) ratios measured in HeLa cells expressing GPDHs from *D. salina, C. reinhardtii, S. arctica* and *D. viridis.* Intracellular levels of DHAP (**g**) and Gro3P (**h**) in HeLa cells expressing GPDHs from *D. salina, C. reinhardtii, S. arctica* and *D. viridis.* Luciferase (LUC) expressing HeLa cells were used as controls in (e-h). _98_*Ds*GPDH_701_, _87_*Cr*GPDH_705_, _100_*Dv*GPDH_701_ and _55_*Sa*GPDH_663_ variants with removed chloroplast targeting sequences and added a C-terminal FLAG tag (See Supplementary Figures S1-S2) were expressed in (c-h). Values are mean ± s.d.; n = 4 in (e, f), n = 3 (g, h) biologically independent samples. Statistically significant differences were calculated by using a one-way ANOVA followed by uncorrected Fisher′s least significant difference test. NS, no significant difference. For growth curves in (d), error bars represent mean ± s.d.; n = 6 biologically independent samples.

NADH-reductive stress is an emerging hallmark of diverse human pathologies, including primary mitochondrial diseases, cancer, cardiac ischemia/reperfusion injury, insulin resistance, fatty liver and dyslipidemia^4–11^. Gro3P accumulation has been identified as a key metabolic feature under NADH-reductive stress, whether induced by the expression of our recently developed genetic tool *Ec*STH (a soluble transhydrogenase that deposits electrons on NAD^+^ using reducing equivalents of NADPH), mitochondrial ETC dysfunction or hypoxia, or by failed pyruvate to lactate conversion via disrupted LDH^12–15^. Gro3P biosynthesis represents a conserved mechanism for NAD^+^ recycling, which protects organisms ranging from yeast and *C. elegans* to mice from NADH-reductive stress under impaired ETC^3,16,17^. However, the accumulation of Gro3P can also have detrimental effects. In human fibroblasts, Gro3P accumulation triggers cellular senescence via excessive lipid accumulation^18^. In the mouse liver, NADH-reductive stress, whether induced by ethanol supplementation or expression of *Ec*STH, activates the transcription factor ChREBP (carbohydrate response element-binding protein), leading to transcription of metabolic programs associated with fatty liver^11^. Interestingly, Gro3P accumulation caused by loss of solute carrier transporter SLC25A13, which leads to Citrin deficiency, was found to directly activate the transcription factor ChREBP to induce expression of fibroblast growth factor 21 (FGF21), which modulates food and alcohol preferences in the brain and contributes to metabolic dysfunction-associated steatotic liver disease (MASLD)[formerly known as non-alcoholic fatty liver disease (NAFLD)]^19^. These findings position elevated NADH/NAD^+^ and Gro3P accumulation as central drivers of ChREBP-mediated metabolic dysfunction in fatty liver disease^11,19^.

Only recently mammalian glycerol-3-phosphate phosphatase (G3PP, encoded by gene *Pgp*) was identified, an enzyme which cleaves Gro3P to glycerol and Pi (**Figure 1a**)^20^. It was postulated that G3PP forms a “glycerol shunt” which re-routes metabolism from accumulation of lipids, especially in organs with high G3PP expression such as heart and skeletal muscle^20^. Moreover, it was demonstrated that modulation of hepatic glycerol shut by G3PP overexpression led to reduced hepatic glucose production and plasma triglycerides levels^20^. In subsequent studies it was also shown that glycerol shunt acts as glucose detoxification pathway in the liver by preventing excessive fat storage^21^, in pancreatic β-cells it controls insulin secretion^22^. Moreover, this mechanism is evolutionary conserved as in *C. elegans* worms the overexpression PGPH-2 (a homolog of G3PP in *C. elegans*) led to decreased fat levels and mimicked the beneficial effects of calorie restriction^23^.

In summary, developing strategies to mitigate NADH-reductive stress and Gro3P accumulation could provide critical insights into the mechanisms underlying metabolic diseases and inform potential therapeutic interventions. At the same time, modulation of endogenous mammalian enzymes of the Gro3P shuttle and the glycerol shunt might not be feasible due to lack of control of the stoichiometry of all reactions involved. Moreover, we were looking to perform experiments in cells when both endogenous Gro3P shuttle and glycerol shunt are intact.

With all that in mind, we turned our attention to di-domain glycerol-3-phosphate dehydrogenases (GPDHs) which were recently identified in several algae^23–25^. These enzymes represent a natural fusion between an N-terminal glycerol-3-phosphate phosphatase (G3PP) and a C-terminal NAD-dependent glycerol-3-phosphate dehydrogenase domain (GPDH) (**Figure 1b**). Therefore, unlike typical mammalian glycerol-3-phosphate dehydrogenases, these fusion di-domain GPDHs not only catalyze NAD-dependent DHAP to Gro3P interconversion but also convert Gro3P to glycerol and inorganic phosphate (Pi) (**Figure 1b**). We reasoned that when expressed in mammalian cells, these enzymes can be used to relieve metabolic consequences of the NADH-reductive stress as they provide simultaneous NAD^+^ recycling and efficient Gro3P clearance (**Figure 1b**). Another advantage of using an enzyme from a lower organism is that it is not subjected to posttranslational or other forms of metabolic regulation compared to its mammalian counterpart^26^. Here, we report the development of a novel genetically encoded tool based on heterologous expression of a di-domain glycerol-3-phosphate dehydrogenase from *Chlamydomonas reinhardtii* (*Cr*GPDH). We demonstrated that *Cr*GPDH relieves metabolic consequences of the NADH-reductive stress under hypoxia or ETC inhibition at either respiratory complex I or III. In addition, we were able to show that *Cr*GPDH expression in clear cell renal cell carcinoma (ccRCC) cells 786-O and Caki-1 reduced triglycerides (TGs) levels. In summary, we anticipate that our new reagent will be instrumental in cellular, organ or whole animal studies aimed at relieving consequences of various metabolic disorders ranging from hypoxia induced injury to dysregulated lipid metabolism in various pathological conditions, including MASLD.

## RESULTS

### Screening di-domain glycerol-3-phosphate dehydrogenases (GPDHs) for their ability to modulate cellular DHAP/Gro3P levels and NAD^+^ recycling in mammalian cells

To facilitate expression in HeLa cells under a doxycycline (Dox)-inducible promoter, constructs encoding four di-domain glycerol-3-phosphate dehydrogenases (GPDHs) from algae (*Dunaliella viridis*, *Dunaliella salina*, *Chlamydomonas reinhardtii* and *Sphaeroforma arctica*) were engineered through *H. sapiens* codon optimization, deletion of predicted chloroplast targeting sequences and incorporation of a C-terminal epitope FLAG tag based on biochemical and structural studies on di-domain glycerol-3-phosphate dehydrogenase from *D. salina* and AlphaFold modeling of all four algae-derived enzymes (**Figure 1b-c**, **Supplementary Figure S1**, **Supplementary Figure S2a-d**)^24^. Expression of GPDH constructs from *D. salina* and *S. arctica* slowed proliferation of HeLa cells while proliferation of cells expressing *C. reinhardtii* and *D. viridis* was not affected (**Figure 1d**, **Supplementary Figure S3a**). We note that all experiments monitoring cellular proliferation were performed in pyruvate-free DMEM supplemented with dialyzed fetal bovine serum (DMEM^+dFBS^). This was done to prevent masking of redox-linked metabolic effects due to pyruvate and other metabolites present in non-dialyzed FBS. We observed that out of four GPDH constructs expressed in HeLa cells, only *C. reinhardtii* variant (*Cr*GPDH) decreased the total cellular NADH/NAD^+^ ratio while its expression did not affect the total cellular NADPH/NADP^+^ ratio (**Figure 1e-f**, **Supplementary Figure S3b-g**). Moreover, expression of *Cr*GPDH was the only GPDH construct which led to a robust decrease in cellular levels of both DHAP and Gro3P (**Figure 1g-h**, **Supplementary Figure S3h-i**). Because *Cr*GPDH expression did not impact proliferation of HeLa cells and robustly decreased both DHAP and Gro3P levels and the NADH/NAD^+^ ratio, we used the *Cr*GPDH construct in the rest of the experiments in this study.

### Biochemical properties of recombinant *Cr*GPDH

To confirm substrate specificity towards NADH and the final products of the catalyzed reaction, we expressed and purified the same _87_*Cr*GPDH_705_-FLAG construct we used in our initial screen in HeLa cells in *E. coli* (**Figure 2a**). The purified _87_*Cr*GPDH_705_-FLAG variant eluted on size-exclusion chromatography as 301 ± 11 kDa protein, suggesting it is a tetramer in solution (**Supplementary Figure S4**). We directly confirmed that the products of the *Cr*GPDH catalyzed reaction are glycerol and Pi, and that this reaction is not detected in the presence of NADPH or the absence of MgCl_2_, as Mg^2+^ is the cofactor for the phosphatase reaction (**Figure 2b-d**). In parallel, we cloned and purified _57_*Cr*GPDH_705_-FLAG variant with an additional 30 amino acids from the unstructured region at the N-terminus to eliminate the possibility that the GPP domain in the construct we used in our initial cellular studies was truncated due to potentially incorrectly predicted boundaries of the chloroplast targeting sequence (CTS) (**Figure 2a**, **Supplementary Figure S1**, **Supplementary Figure S2b**, **Supplementary Figure S4**). Both glycerol and Pi formation were present for both _57_*Cr*GPDH_705_-FLAG and _87_*Cr*GPDH_705_-FLAG variants while no traces of Gro3P were detected in our enzymatic assays (**Figure 2c-d**). Moreover, Gro3P was fully consumed but only in the presence of Mg^2+^ when it was used in assays with recombinant enzymes (**Figure 2d**). This further confirmed that both _57_*Cr*GPDH_705_-FLAG and _87_*Cr*GPDH_705_-FLAG variants possess fully active N-terminal GPP domain (which converts Gro3P to glycerol and Pi). In the rest of the study, we refer to the _87_*Cr*GPDH_705_-FLAG variant as *Cr*GPDH. Using a continuous assay, we determined *K*_M_’s of recombinant *Cr*GPDH as 45 ± 3 μM for NADH and 820 ± 127 μM for DHAP, while *k*_cat_’s were 147 ± 10 and 248 ± 36 s^−1^, respectively (**Figure 2e-f**, **Table 1**). In summary, our biochemical characterization confirmed that *Cr*GPDH is a strictly NADH-specific di-domain GPDH and its reaction products are NAD^+^, glycerol and P_i_.

**Figure 2:**
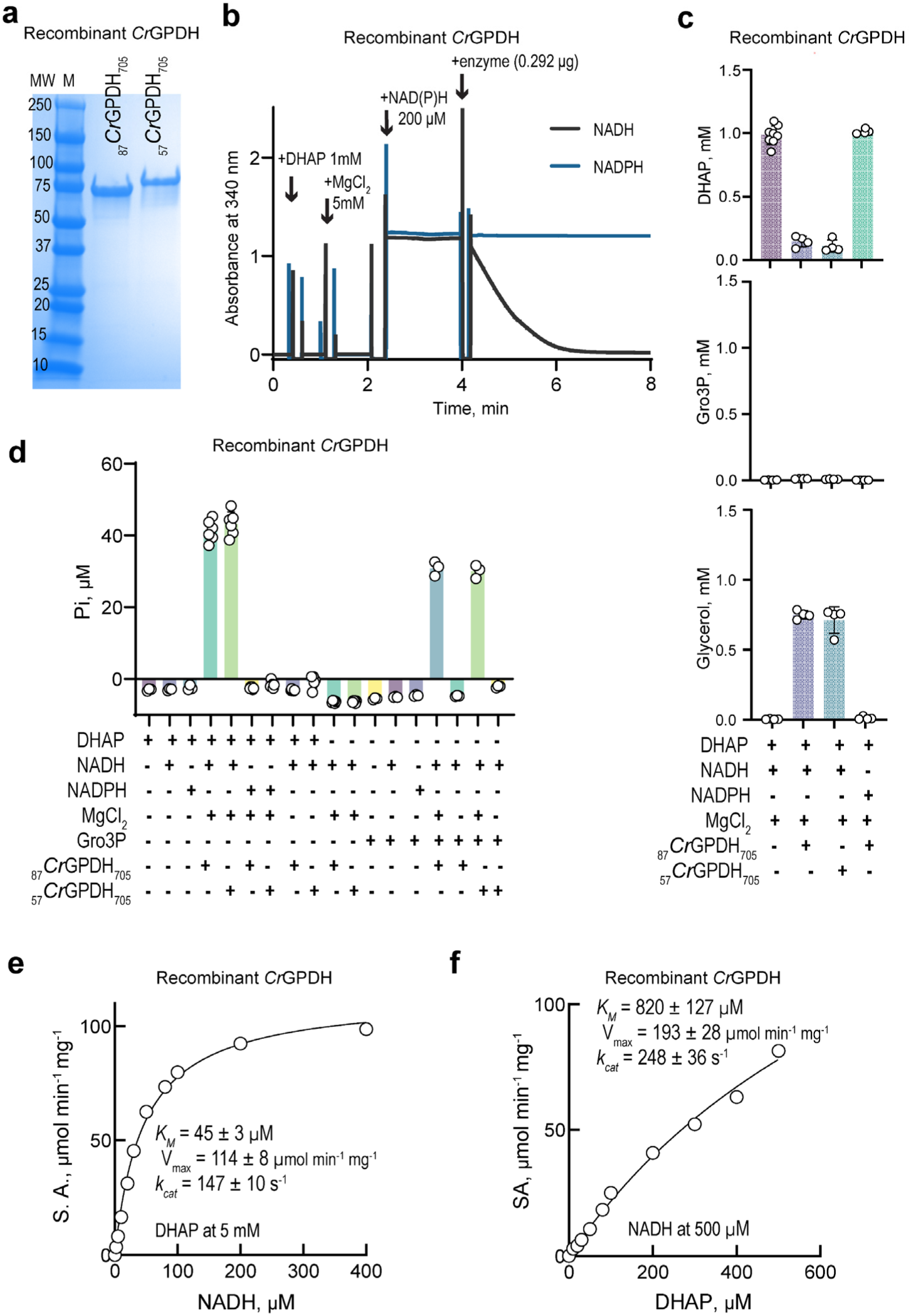
Biochemical properties of GPDH from *C. reinhardtii*. **(a)** SDS-PAGE of purified recombinant _87_*Cr*GPDH_705_ and _57_*Cr*GPDH_705_ variants. A representative SDS-PAGE gel is shown. **(b)** Representative time courses of the reaction catalyzed by recombinant _87_*Cr*GPDH_705_ variant in the presence of 200 µM NADH or NADPH. **(c)** Detection of DHAP, Gro3P and glycerol by GC-MS in reactions with recombinant _87_*Cr*GPDH_705_ and _57_*Cr*GPDH_705_ variants. Reaction mixture in (c) contained 1 mM DHAP, 1 mM NADH or NADPH and _87_*Cr*GPDH_705_ or _57_*Cr*GPDH_705_ enzymes and other components as indicated. Ten minutes after incubation at 37 °C, reactions were quenched with equal volume of 80%/20% methanol/H_2_O containing 100 µM norvaline, derivatized and subjected to GC-MS analysis. **(d)** Pi production by recombinant _87_*Cr*GPDH_705_ and _57_*Cr*GPDH_705_ variants as determined by the malachite green assay in the presence of 50 µM DHAP or Gro3P and other components in the reaction mixture as indicated. Michaelis-Menten analysis of the reaction catalyzed by _87_*Cr*GPDH_705_ with NADH (**e**) or DHAP (**f**) as a varying substrate. In (e) DHAP was fixed at 5 mM and in (f) NADH was fixed at 500 µM. Reported values in (e, f) for V_max_, *k*_cat_ and *K*_M_ are taken from Table 1. Details of enzymatic assays in (b-f) are described under Methods. Values are mean ± s.d.; n = 8, 4, 4, 4 in (c), n = 3, 3, 3, 6, 6, 6, 6, 6, 6, 6, 6, 3, 3, 3, 3, 3, 3, 3 in (d), n = 2 in (e, f) biologically independent samples.

**Table 1:** Steady-state kinetic parameters of *Cr*GPDH.

| Substrate pair<sup>a</sup> | Varying Substrate | $K_m$<br>($\mu\text{M}$) | $V_{\text{max}}$<br>($\mu\text{mol min}^{-1}$<br>$\text{mg}^{-1}$) | $k_{\text{cat}}$<br>($\text{s}^{-1}$) | $k_{\text{cat}}/K_m$<br>($\text{s}^{-1} \text{M}^{-1}$) |
| --- | --- | --- | --- | --- | --- |
| NADH + DHAP | NADH | $45 \pm 3$ | $114 \pm 8$ | $147 \pm 10$ | $(3.2 \pm 0.3) \times 10^6$ |
| NADH + DHAP | DHAP | $820 \pm 127$ | $193 \pm 28$ | $248 \pm 36$ | $(0.30 \pm 0.06) \times 10^6$ |
| NADPH<br>+<br>DHAP |  | no reaction<sup>b</sup> |  |  |  |

<sup>a</sup>For all substrate pairs, activity was measured using *87Cr*GPDH<sub>705</sub> variant by following the absorbance at 340 nm (the dehydrogenase reaction). When NADH was varying, substrate DHAP was fixed at 5 mM; when DHAP was varying, substrate NADH was fixed at 500 $\mu\text{M}$ . <sup>b</sup>No activity was detected with NADPH. Details of enzymatic assays are described under Methods.

### Metabolic features of *Cr*GPDH expression in mammalian cells

In addition to the decreased levels of DHAP and Gro3P when expressed in HeLa cells, *Cr*GPDH affected a few metabolites of central carbon metabolism (**Figure 3a-h**). The “KEGG” enrichment terms analysis revealed that purine metabolism and pentose phosphate pathway (PPP) are the most affected metabolic pathways with *Cr*GPDH expression (**Figure 3b**). Interestingly, except for increased pyruvate accumulation and DHAP and Gro3P consumption, metabolites comprising glycolysis remained largely unaffected by *Cr*GPDH expression (**Figure 3d**). Notably, metabolites involved in PPP, purine and pyrimidine metabolism were slightly decreased, whereas TCA cycle intermediates increased in HeLa cells expressing *Cr*GPDH compared to luciferase (LUC) control (**Figure 3e-h**). Both pyruvate and TCA cycle intermediates accumulation is consistent with the NAD^+^ recycling activity of *Cr*GPDH, which is similar to the NAD^+^ recycling activity of a water-forming NADH oxidase *Lb*NOX^16,27^. Interestingly, the metabolites from PPP, purine, and pyrimidine metabolism that decreased under *Cr*GPDH expression all contain sugar phosphates. This suggests that an excess of inorganic phosphate (Pi) produced by *Cr*GPDH could affect sugar phosphate biosynthesis and signaling. Of note, expression of *Cr*GPDH did not impact cellular oxygen consumption rate (OCR) and only slightly decreased extracellular acidification rate (ECAR), which is a proxy for glycolysis (**Figure 3i-j**, **Supplementary Figure S5a-d**).

**Figure 3:**
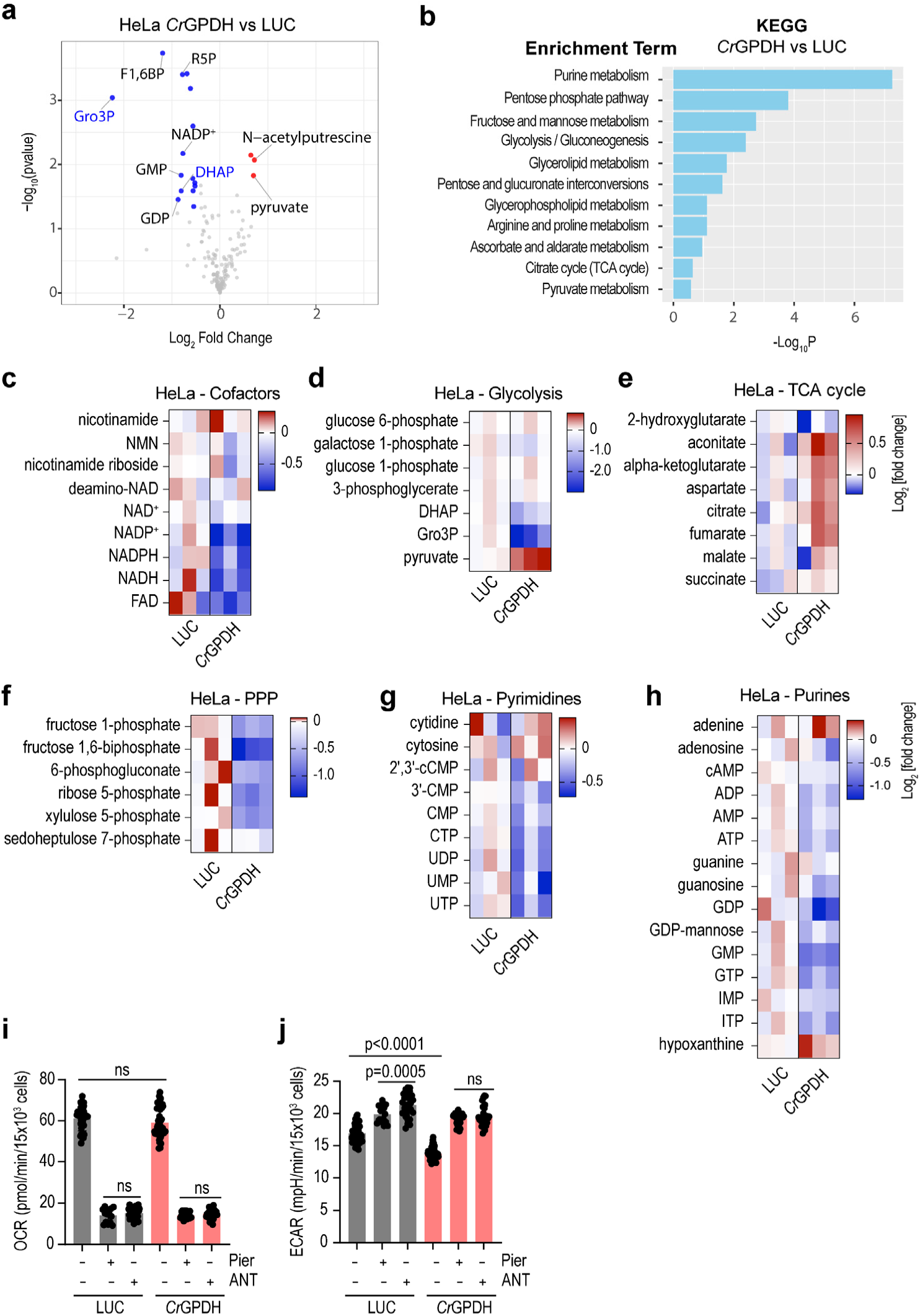
Metabolic features of HeLa cells expressing *Cr*GPDH. (**a**) Volcano plots for targeted metabolomics of HeLa cells expressing *Cr*GPDH when compared to LUC. The statistical significance (accumulated metabolites shown in red dots, decreased metabolites shown in blue dots) represents p value cutoff = 0.05, fold change cutoff = 0.5, gray dots represent statistically not significant changes. DHAP, dihydroxyacetone phosphate; Gro3P, glycerol-3-phosphate; F1,6BP, fructose-1,6-biphosphate; R5P, ribose-5-phosphate. (**b**) Kyoto Encyclopedia of Genes and Genomes (KEGG) pathway enrichment analysis for metabolites in (a) which were significantly changed. Heatmaps of the most impacted intracellular cofactors (**c**), glycolysis (**d**), TCA cycle (**e**), PPP (**f**), pyrimidines (**g**) purines (**h**) metabolites in HeLa cells expressing *Cr*GPDH and LUC. Oxygen consumption rate (OCR) (**i**) and extracellular acidification rate (ECAR) (**j**) of HeLa cells expressing *Cr*GPDH before and after separate additions of 1 µM piericidin A (Pier) or 1 µM antimycin A (ANT) measured in pyruvate free HEPES/DMEM^+dFBS^ media. Values are mean ± s.d.; n = 45, 15, 30, 55, 25, 30 in (i), n = 50, 20, 30, 60, 30, 30 in (j) biologically independent samples. The statistical significance indicated for (i-j) represents a One-Way ANOVA followed by Šídák multiple comparison test. LUC was used as controls in (a-j). In heatmaps (c-h), each column represents a biologically independent sample.

### *Cr*GPDH expression bypasses growth arrest mediated by inhibition of mitochondrial ETC or hypoxia

ETC inhibition leads to a growth arrest and, depending on the exact complex within the ETC that is inhibited, may require pyruvate and uridine in the culture medium to allow mammalian cells to proliferate ^28^. We previously demonstrated that ectopic expression of *Lb*NOX promotes NAD^+^ recycling and can be used instead of pyruvate or other extracellular electron acceptors such as α-ketobutyrate (AKB) or oxaloacetate (OAA) to normalize NADH/NAD^+^ ratio and rescue ETC-mediated growth arrest^16^. To evaluate whether *Cr*GPDH has a similar function, we evaluated the proliferation of HeLa cells expressing *Cr*GPDH under various growth conditions in DMEM^+dFBS^. We observed that *Cr*GPDH expression completely rescued growth arrest associated with mitochondrial complex I inhibition and partially rescued growth arrest associated with mitochondrial complex III inhibition in HeLa cells grown in DMEM^+dFBS^ medium in the absence of pyruvate and uridine (**Figure 4a-c**). This clearly demonstrated that *Cr*GPDH expression promotes NAD^+^ recycling in cells with inhibited mitochondrial complexes I or III and allows cells to proliferate.

**Figure 4:**
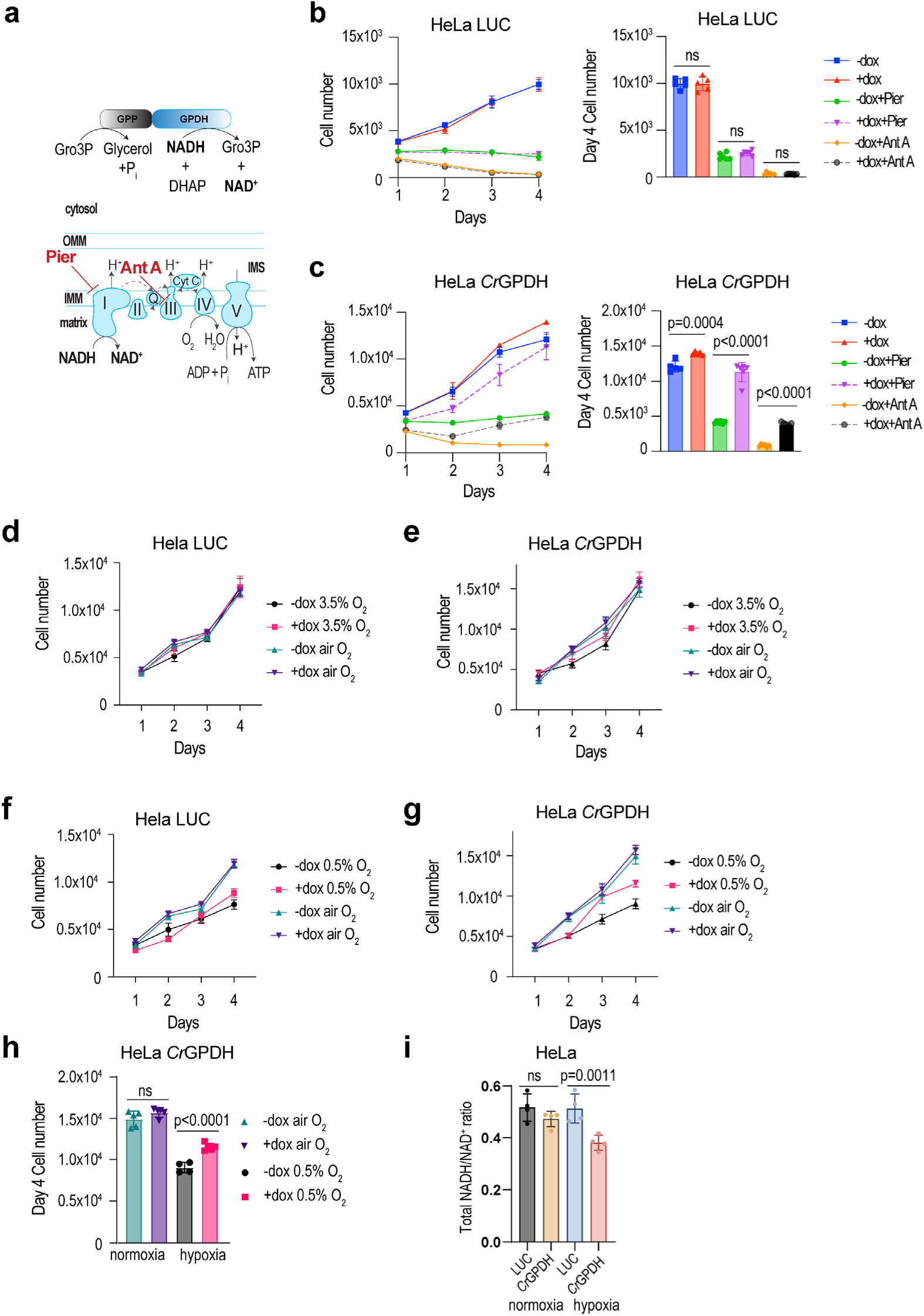
*Cr*GPDH expression allows partial alleviation of growth arrest of HeLa cells with inhibited ETC or under hypoxia. (**a**) Schematic of *Cr*GPDH catalyzed reaction expressed in the cytoplasm and mitochondrial ETC complex I inhibition with piericidin A and complex III inhibition with antimycin A. Effect of LUC (**b**) and *Cr*GPDH (**c**) expression on proliferation of HeLa cells grown in DMEM^+dFBS^ in the absence of pyruvate and uridine with inhibited complex I (1 µM piericidin) or inhibited complex III (1 µM antimycin). In (b-c), cell numbers at day 4 are shown as bar graphs for all conditions specified. Proliferation of HeLa cells expressing LUC or *Cr*GPDH at 3.5% O_2_ hypoxia (**d-e**) and 0.5% O_2_ hypoxia (**f-g**). Cell numbers at day 4 of HeLa cells with and without *Cr*GPDH expression grown in room air (normoxia) or under 0.5% O_2_ (**h**). The total cellular NADH/NAD^+^ ratio measured in HeLa cells expressing LUC or *Cr*GPDH under normoxia or 0.5% O_2_ hypoxia (**i**). For growth curves in (b, c, d-g), error bars represent mean ± s.d.; n = 6 biologically independent samples. Values are mean ± s.d.; n = 5, 5, 6, 6, 6, 6 in (b), n = 5, 5, 6, 6, 6, 6 in (c), n = 5, 5, 4, 5 in (h), n = 4 in (i) biologically independent samples. The statistical significance indicated for (b-c) represents a One-Way ANOVA followed by Šídák multiple comparison test. The statistical significance indicated for (b-c) represents a One-Way ANOVA followed by Šídák multiple comparison test. The statistical significance indicated for (h-i) represents a One-Way ANOVA followed by uncorrected Fisher′s least significant difference test. NS, no significant difference.

Limited availability of oxygen as a final electron acceptor under hypoxia conditions also contributes to low ETC activity, elevated NADH/NAD^+^ ratio and growth arrest^14,29,30^. Unlike *Lb*NOX, *Cr*GPDH does not need oxygen as a terminal electron acceptor, so we decided to test whether NAD^+^ recycling by *Cr*GPDH could rescue hypoxia-induced growth defects. We monitored the growth of HeLa cells expressing LUC or *Cr*GPDH under multiple oxygen concentrations. At 3.5% oxygen, no difference was observed between 21% (normoxic air) and hypoxia in both LUC and *Cr*GPDH expressing HeLa cells (**Figure 4d-e**). We next observed that at 0.5% O_2_ hypoxia, proliferation of HeLa cells expressing LUC was slower compared to that at 21% O_2_ but *Cr*GPDH expression allowed substantial growth rescue after two days following induction of *Cr*GPDH expression with Dox (**Figure 4f-h**). Interestingly, under 0.5% O_2_ hypoxia, we observed a clear decrease in NADH/NAD^+^ ratio in cells expressing *Cr*GPDH compared to LUC expressing controls, suggesting a robust activity of *Cr*GPDH under these conditions (**Figure 4i**). Taken together, *Cr*GPDH expression promotes NAD^+^ regeneration and supports proliferation under normoxia with pharmacologic ETC inhibition or under 0.5% O_2_ hypoxia. Our observations align with previous studies demonstrating that NAD^+^ regeneration through Gro3P biosynthesis can be an important endogenous mechanism to bypass ETC inhibition^3^. In our system, *Cr*GPDH further converts Gro3P to glycerol and Pi, which prevents Gro3P accumulation and allows uninterrupted NAD^+^ recycling (because mGPDH is coenzyme Q linked, diminished electron flow via ETC decreases the activity of endogenous Gro3P shuttle and we previously observed elevated levels of both DHAP and Gro3P under conditions of NADH-reductive stress) (**Figure 1a**)^15^.

### Transcriptomic features of *Cr*GPDH expression in mammalian cells

To further understand the effect of *Cr*GPDH expression on metabolism, we performed a bulk RNA sequencing (RNA-seq) analysis of HeLa cells with *Cr*GPDH expression (**Figure 5a-b**). Consistent with the metabolic profiling, HeLa cells with *Cr*GPDH expression showed modest changes in gene expression, with only 19 upregulated and 45 downregulated differentially expressed genes (DEGs) (**Figure 5a**). Furthermore, Gene Ontology (GO) term analysis revealed that the DEG sets associated with *Cr*GPDH expression were related to cell differentiation, extracellular matrix proteins, and protein phosphatase activity (**Figure 5b**). The Cnetplots showed a strong link between the upregulation of IL6, HES1, RORA, and MMP11 genes and “fat cell differentiation” GO terms (**Figure 5c**). It also highlighted the downregulation of DUSP4, DUSP5, and GO terms linked to MAP kinase phosphatase (MKP) activity (**Figure 5d**). Interestingly, DUSP4 and DUSP5 are part of the dual-specificity phosphatases (DUSPs) family. They inactivate MAP kinases by dephosphorylating threonine/tyrosine residues in the T-X-Y motif of the kinase activation loop^31^. This suggests that HeLa cells expressing *Cr*GPDH downregulate DUSPs to promote MAP kinase activity. This likely reflects the intrinsic adaptation to the accumulation of inorganic phosphate (Pi) produced by *Cr*GPDH. We also observed the upregulation of the SLC38A3 gene which is believed to encode a symporter for glutamine, asparagine and histidine sodium ions that is coupled to an H^+^ antiporter activity^32–34^. Taken together, *Cr*GPDH expression in HeLa cells impacts cellular programming linked to proliferation, differentiation and MAPK signaling.

**Figure 5:**
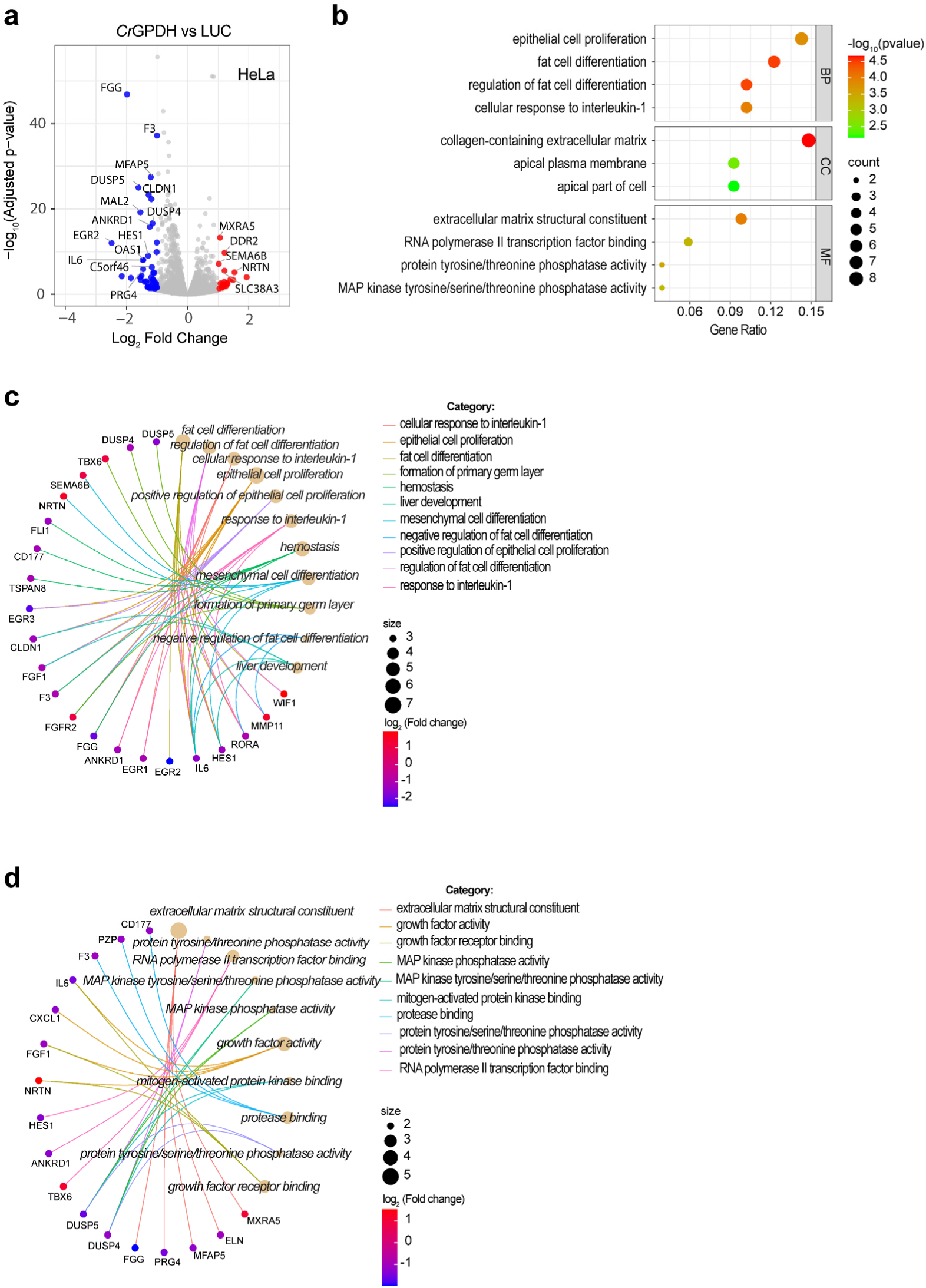
Analysis of transcriptomic features of *Cr*GPDH expression in HeLa cells. (**a**) Volcano plots that represent the log2 fold change (x axis) and adjusted p value for significance (y axis) of *Cr*GPDH vs LUC expressing HeLa cells. Genes significantly different in expression at false discovery rate (FDR) of 5% are indicated in red (upregulated genes, log_2_ fold change above 1) or blue (downregulated genes, log_2_ fold change below −1). Gray dots represent genes without significant changes. (**b**) Gene ontology (GO) terms analysis of significant genes that are differentially changed between *Cr*GPDH and LUC. BP: biological process; CC, cellular component; MF: molecular function. (c, d) The Cnet plots for the subset of genes that correlate with GO terms BP in (**c**) and MF in (**d**).

### *Cr*GPDH expression increases fatty acid β-oxidation in HeLa cells

Because DHAP acts as a precursor for Gro3P, and the latter is an essential intermediate in the biosynthesis of phospholipids and triacylglycerols (TGs), we performed lipidomic profiling of HeLa cells expressing *Cr*GPDH (**Figure 6a-b**). Lipidomic profiling revealed a substantial decrease in acylcarnitine levels without affecting other lipid classes in HeLa cells expressing *Cr*GPDH when compared to LUC control (**Figure 6a-b**). Notably, the detected acylcarnitines (18:1), (16:0), (14:0) and (16:1) are all long-chain acylcarnitines (C13–C20) (**Figure 6b**), which are produced by the carnitine shuttle to transport long chain fatty acids into mitochondria for fatty acid β-oxidation (FAO). Accumulated long-chain acylcarnitines are diagnostic markers for inherited FAO disorders^35,36^. Thus, it is reasonable to assume that the decreased level of long chain acylcarnitines in HeLa cells with *Cr*GPDH expression is due to activation of mitochondrial FAO. Interestingly, overlapping *Cr*GPDH-related DEGs with the human mitochondrial gene database, MitoCarta3.0^37^, revealed upregulation of multiple genes encoding enzymes linked to FAO (**Figure 6c-d**). For example, we observed upregulation of ACSF2 (medium-chain fatty acid-CoA ligase), an enzyme that was previously shown to be localized in the mitochondria, where it activates medium-chain fatty acids by converting them to coenzyme A thioesters (medium-chain fatty acids do not require the carnitine shuttle to enter mitochondria)^38,39^. Two enzymes that are part of the FAO pathway, short chain acyl-CoA dehydrogenase (encoded by ACADS) and enoyl-CoA hydratase/3-hydroxyacyl CoA dehydrogenase (encoded by EHHADH) were also upregulated in *Cr*GPDH expressing cells. Moreover, genes HMGCL (cleaves 3-hydroxy-3-methylglutaryl-CoA into acetyl-CoA and acetoacetate) and CLYBL (cleaves citramalyl-CoA into acetyl-CoA and pyruvate)^40^ are also upregulated with *Cr*GPDH expression. Interestingly, the SLC25A42 gene, which encodes the mitochondrial transporter for CoASH, was also upregulated in HeLa cells with *Cr*GPDH expression. We also observed upregulation of CPT1C, an isoform of carnitine palmitoyl transferase I, which was shown to be expressed in endoplasmic reticulum of adult mouse neurons^41^. In summary, our findings suggest that *Cr*GPDH expression promotes increased degradation of fat resources for energy metabolism (acetyl-CoA produced in β-oxidation and by HMGCL and CLYBL can be directly fed into the TCA cycle), and diverting Gro3P into glycerol does not affect downstream lipogenesis in HeLa cells.

**Figure 6:**
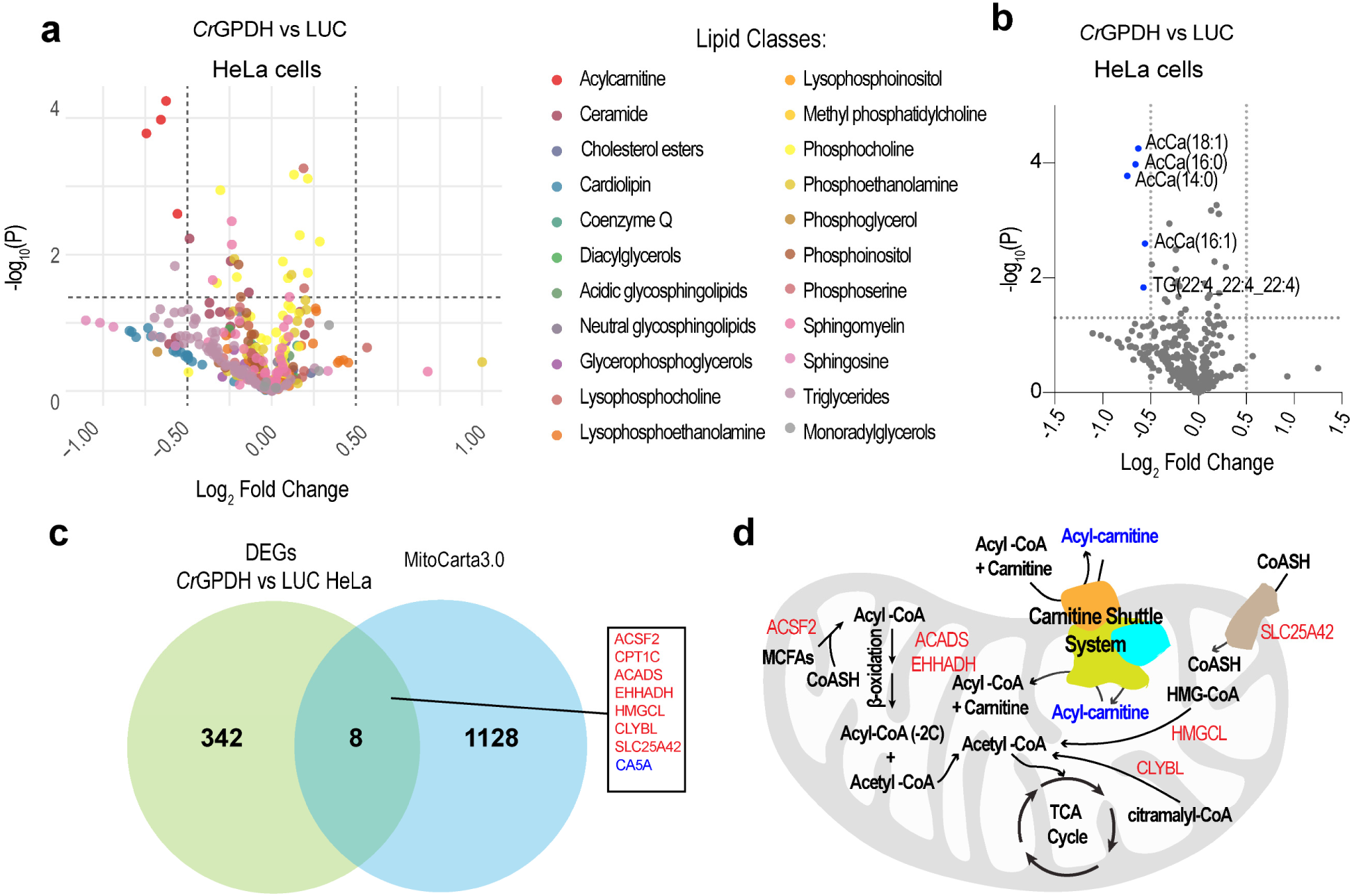
*Cr*GPDH expression increases fatty acid β-oxidation in HeLa cells. (**a-b**) Volcano plots showing the log_2_ fold change (x axis) and p value (y axis) of lipidomic profiling of HeLa cells with *Cr*GPDH expression compared to LUC. Lipids are color-coded by class in (a). Significantly changed lipids (p value cutoff = 0.05, fold change cutoff = 0.5) are highlighted in blue, while gray dots represent lipids without significant changes in (b). AcCa: acylcarnitine, TG: triglycerides. (**c**) A Venn diagram showing the overlap between the 350 *Cr*GPDH vs LUC differentially expressed genes (DEGs) and the 1136 genes encoding the human mitochondrial proteome from MitoCarta3.0. The cutoff for DEGs is a log_2_ fold change > 0.5. Upregulated overlapping genes are in red, while downregulated ones are in blue. (**d**) The schematics depicting fatty acid β-oxidation (FAO) and related pathways. Decreased acylcarnitines from (b) are highlighted in blue. Correlated genes in (c) are highlighted in red. MCFAs: medium chain fatty acids.

### Decreased lipogenesis in clear cell renal cell carcinoma (ccRCC) cells expressing *Cr*GPDH

Clear cell renal cell carcinoma (ccRCC) is the predominant type of kidney cancer, characterized by abnormal lipid accumulation^42^. A recent study has shown that lipid synthesis in kidney cancers is supported by diverting the Gro3P shuttle towards an increased Gro3P biosynthesis^43^. To investigate whether *Cr*GPDH could affect this lipid accumulation, we expressed *Cr*GPDH in two widely used ccRCC cell lines, 786-O and Caki-1. Since 786-O cells are sensitive to doxycycline (Dox), we constructed plasmids for constitutive expression of *Cr*GPDH and LUC in 786-O cells. For Caki-1 cells, we used a Dox-inducible system to express *Cr*GPDH and LUC. The robust expression of *Cr*GPDH in both 786-O and Caki-1 cell lines was validated by Western blot (**Figure 7a-b**). Consistent with the results in HeLa cells, both 786-O and Caki-1 cells expressing *Cr*GPDH showed a significant decrease in the total cellular NADH/NAD^+^ ratio without affecting the total cellular NADPH/NADP^+^ ratio (**Figure 7c-d**), while exhibiting normal cell proliferation compared to controls (**Figure 7e-f**). Further lipidomic profiling revealed a dramatic decrease in TGs levels in 786-O cells and a more moderate impact in Caki-1 cells expressing *Cr*GPDH compared to the LUC control (**Figure 7g-h**, **Supplementary Figure S6**, **Supplementary Figure S7**), suggesting that *Cr*GPDH actively clears Gro3P in kidney cancer cells, thereby inhibiting associated lipogenesis (**Figure 8a-b**).

**Figure 7:**
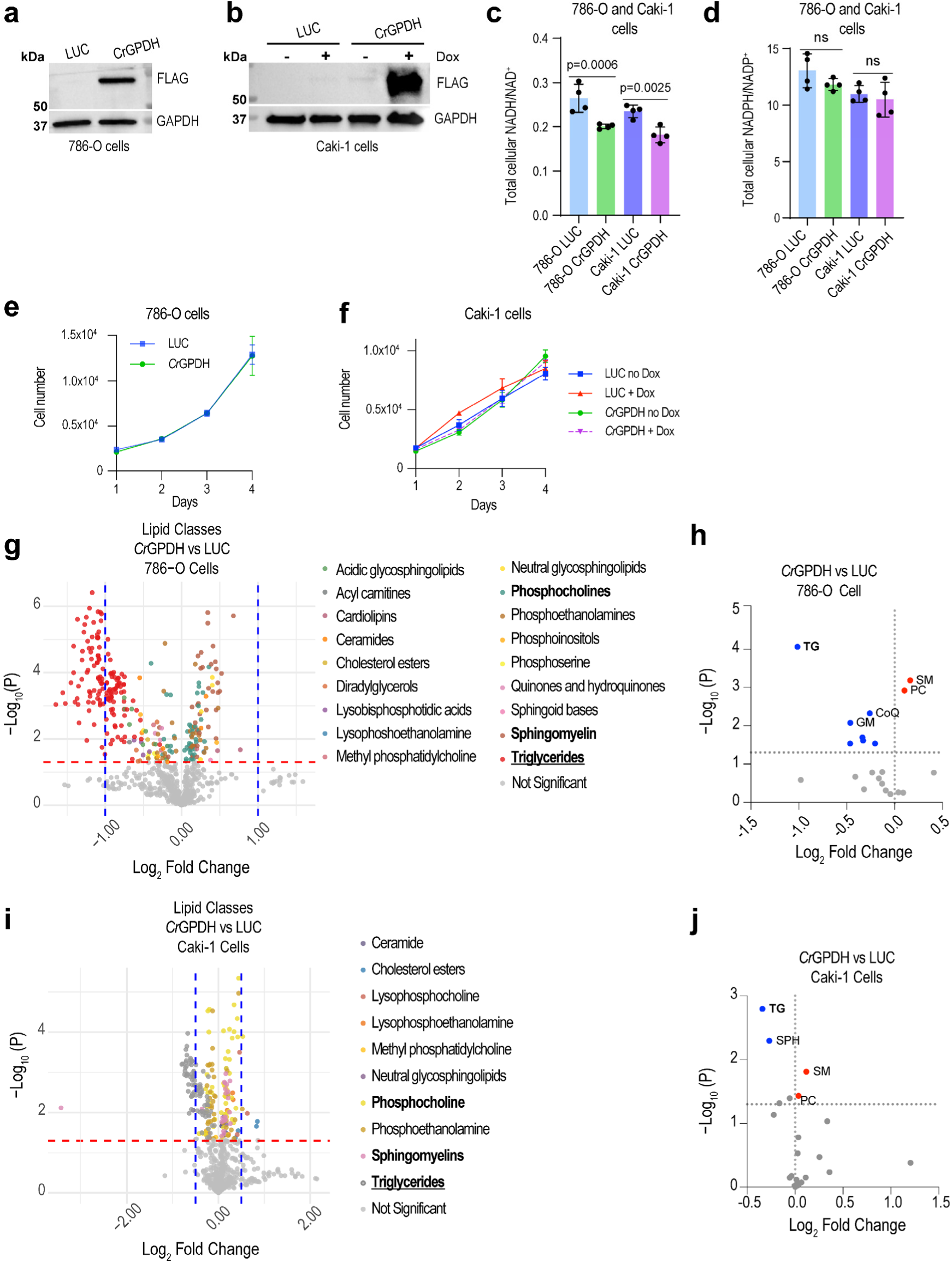
*Cr*GPDH alters lipid profiles of 786-O and Caki-1 clear cell renal cell carcinoma (ccRCC) cell lines. (**a**, **b**) Western blot of 786-O and Caki-1 cells expressing *Cr*GPDH with a FLAG tag and Luciferase (LUC). *Cr*GPDH was expressed in Caki-1 cells under Dox control (24 h after Dox addition). A representative Western blot is shown. The total cellular NADH/NAD^+^ (**c**) and NADPH/NADP^+^ (**d**) ratios measured in 786-O and Caki-1 cells expressing *Cr*GPDH and LUC. (**e**, **f**) The effect of expression of *Cr*GPDH on proliferation of 786-O and Caki-1 cells grown in pyruvate-free RPMI^+dFBS^ and DMEM^+dFBS^, respectively. Volcano plots show the log_2_ fold change (x-axis) and p-value (y-axis) for individual lipid color-coded by class (**g, i**) or summed lipid classes (**h**, **j**) in 786-O and Caki-1 cells expressing *Cr*GPDH. TG: triglycerides, SM: sphingomyelin, PC: phosphocholines, CoQ: coenzyme Q-like molecules, GM: gangliosides of acidic glycosphingolipids, SPH: sphingosine. Values are mean ± s.d.; n = 4 in (c, d) biologically independent samples. Statistically significant differences were calculated by using a One ANOVA followed by followed by uncorrected Fisher′s least significant difference test. NS, no significant difference. For growth curves in (e-f), error bars represent mean ± s.d.; n = 6 biologically independent samples.

**Figure 8:**
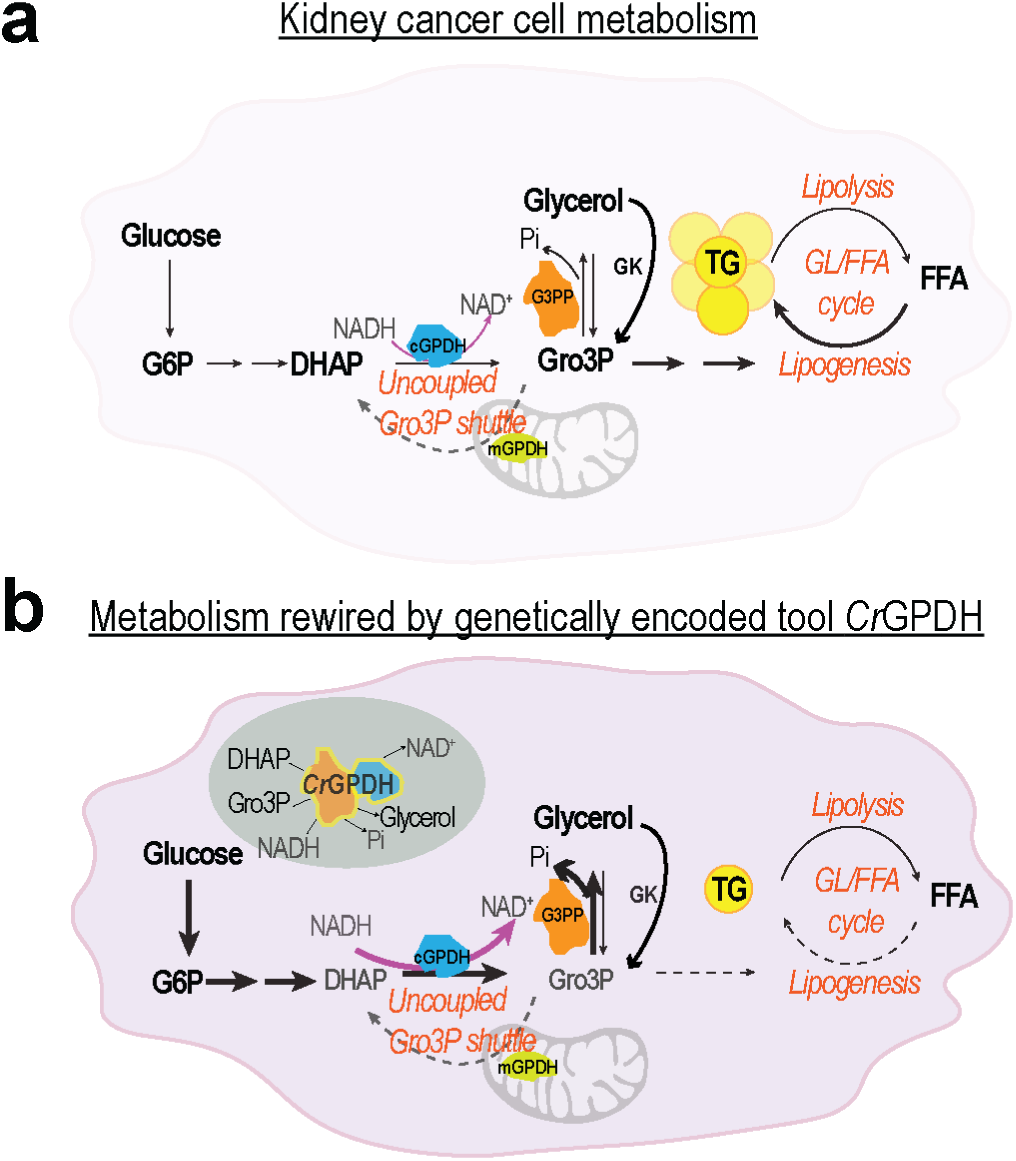
Diagram summarizing metabolic rewiring in clear cell renal cell carcinoma (ccRCC) cells expressing *Cr*GPDH. FFA: free fatty acid; TG: triglycerides; GL/FFA cycle: glycerolipid/free fatty acid cycle; GK: glycerol kinase.

## DISCUSSION

### Elevated cellular levels of oxidized NAD^+^ increased mitochondrial TCA cycle flux and fatty acid β-oxidation

Here, we show that *Cr*GPDH can effectively regenerate NAD^+^ and consume both DHAP and Gro3P in both *in vitro* assays with purified proteins and mammalian cell culture. The physiological adaptation of mammalian cells to a low NADH/NAD^+^ environment restricts the potential side effects of boosting the pro-oxidative shift (a decrease in NADH/NAD^+^ ratio). Unlike the substantial changes in transcriptome and metabolome under NADH-reductive stress, HeLa cells expressing *Cr*GPDH exhibited minor metabolic and transcriptomic changes, which is in line with findings from our previous study where genetically encoded tool *Lb*NOX was used to recycle NAD^+15^ (**Figure 3a-h**, **Figure 5a-d**). We also observed pyruvate accumulation when NAD^+^ was recycled wither either *Lb*NOX^15^ or *Cr*GPDH as in both cases, cells are less dependent on the pyruvate-consuming LDH-catalyzed reaction to normalize NADH/NAD^+^ ratio^16^. We also found that the TCA cycle intermediates are accumulated in HeLa cells expressing *Cr*GPDH (**Figure 3e**), suggesting that more glucose carbons were diverted into the TCA cycle to maintain NADH production, something which was also previously demonstrated for *Lb*NOX^16,27^. We note that because *Cr*GPDH does not require O_2_ as a co-substrate our new genetic tools can be used in various applications in hypoxia as it allows to decrease cellular NADH/NAD^+^ under these conditions (**Figure 4i**). Moreover, our comprehensive multi-omics analysis of cells expressing *Cr*GPDH revealed activation of mitochondrial FAO (**Figure 6c-d**). This is likely due to the increased levels of NAD^+^, which acts as an electron acceptor in both the TCA cycle and mitochondrial FAO. Despite these changes, the bioenergetic features (OCR and ECAR) remained unaffected (**Figure 3i, j**). This suggests that the reducing equivalents from the increased TCA flux and FAO do not enter the mitochondrial ETC but likely instead support the dehydrogenase activity of the *Cr*GPDH tool, which catalyzes the conversion of DHAP into Gro3P by regenerating NAD^+^, forming a positive feedback loop that crosslinks glucose, lipid, and energy metabolism.

### Cellular response to the accumulation of inorganic phosphate (Pi)

We found decreased levels of PPP, purine and pyrimidine metabolites in HeLa cells expressing *Cr*GPDH (**Figure 3f-h**). Notably, multiple metabolites with decreased accumulation under *Cr*GPDH expression were sugar phosphates (**Figure 3c-h**). The decreased sugar phosphates are likely regulated by the inorganic phosphate (Pi) produced by *Cr*GPDH. The expression of DUSPs genes (encoding phosphatases that inactivate MAP kinases) are downregulated in HeLa cells expressing *Cr*GPDH, indicating stimulated MAPK signaling in these cells (**Figure 5d**). Our findings likely reflect key cellular protein post-translational adaptations to the accumulation of inorganic phosphate (Pi) produced by *Cr*GPDH to sense phosphate availability and modify protein activity by phosphorylation. Notably, mammals lower blood phosphate levels by enhancing kidney glycolysis and stimulating the synthesis of Gro3P through the activation of cGPDH^44^. The increased Gro3P then circulates to bone cells to stimulate the bone-derived hormone fibroblast growth factor 23 (FGF-23), which reduces blood phosphate levels by blocking kidney absorption, thus forming a kidney-bone feedback loop to maintain blood phosphate homeostasis^44^. This suggests that the phosphate produced by *Cr*GPDH expression has the potential to activate endogenously expressed cGPDH, forming a feedback loop that potentially maximizes Gro3P synthesis and NAD^+^ recycling and stimulates upstream glycolysis to make more fuels for energy metabolism. However, it is not clear how cGPDH activity is activated by circulating Gro3P and FGF-23. We speculate that GPD1 protein activity is likely activated via protein post-translational modifications, such as phosphorylation by MAPK signaling. However, studies in yeast demonstrate that cGPDH activity is inhibited by phosphorylation at multiple sites, as evidenced by increased activity in mutants lacking these phosphorylation sites^45^. This raises the possibility that phosphorylation-dependent regulation of mammalian cGPDH may differ from yeast. Future studies should investigate how MAPK signaling modulates cGPDH activity under conditions of phosphate excess, particularly through site-specific phosphorylation.

### Benefits from reduced triglycerides (TGs) accumulation

*Cr*GPDH expression reduced triglycerides (TGs) accumulation in two clear cell renal cell carcinoma (ccRCC) cell lines but not in HeLa cells (**Figure 6a-b**, **Figure 7g-j**). This discrepancy is likely due to differences in lipid composition between cancer cell lines. In HeLa cells, TGs constitute only a minor fraction of the total lipid profile^46^, whereas in kidney lipid extracts, TGs represent the most abundant lipid class^43^. Due to the limited total amount of TGs in HeLa cells, the minor TGs loss by *Cr*GPDH expression in HeLa cells can be easily compensated by *de novo* lipogenesis from acetyl-CoA. This is likely due to robust activation of multiple metabolic pathways that generate acetyl-CoA in HeLa cells, including FAO (**Figure 6d**). This discrepancy can also be regulated by the uncoupled Gro3P shuttle mechanism in kidney cancer cells which redirects carbon flux into lipid synthesis via increased cytosolic to mitochondria GPDH (cGPDH/mGPDH) ratio^43^. Moreover, uncoupled Gro3P shuttle was also observed in humans and mice with diabetes and obesity, leading to excessive lipid accumulation and cardiomyopathy^47^. These studies suggest that lipid accumulation by an uncoupled Gro3P shuttle contributes to kidney cancers and heart dysfunction in metabolic disorders. Notably, it was shown that cell proliferation of kidney cancers depends on lipid synthesis but not NAD^+^ recycling via the uncoupled Gro3P shuttle^43^. Conversely, we show that decreased levels of TGs in ccRCC cell lines expressing *Cr*GPDH do not affect proliferation (**Figure 7e-f**). This finding contrasts with several other studies that revealed distinct functions of cytosolic GPDH (cGPDH) in cancer metabolism^48–51^. Therefore, future work is required to systematically study Gro3P metabolism in various cancer types.

Accumulation of TGs in normal tissues can lead to conditions such as MASLD (formerly known as NAFLD), insulin resistance and diabetes^52,53^. With the capability of decreasing TGs, our novel *Cr*GPDH genetic tool provides a new opportunity to alleviate disrupted lipid metabolism in these pathologies. Our new reagent *Cr*GPDH allows to reliably downregulate TGs levels by diverting glycolytic Gro3P into glycerol similarly to a “stand-alone” glycerol shunt catalyzed by G3PP (**Figure 1a-b**, **Figure 8a-b**) ^20,21^. We also note that cGPDH activity can be activated by ergothioneine, a diet-derived, atypical amino acid, and this effect was linked to an increased healthspan and lifespan in aged rats and *C. elegans*^54^. In summary, our findings make di-domain *Cr*GPDH a promising tool for future applications aimed at tackling multiple pathologies linked to impaired lipid metabolism.

## Materials and Methods

### Cell Culture

HeLa cells were cultured in DMEM^+FBS^ [DMEM without pyruvate and glucose (ThermoFisher, 11966025) supplemented with 25 mM glucose, 10% non-dialyzed FBS (Sigma, F2442) and 1% penicillin/streptomycin] at 37°C in 5% CO_2_. Lenti HeLa Tet-One-Puro cell lines were cultured in DMEM^+FBS^ in the presence of 1 µg/mL puromycin. All experiments with lenti HeLa Tet-One-Puro cell lines were performed in antibiotics-free DMEM^+dFBS^ [DMEM without pyruvate and glucose supplemented with 25 mM glucose and 10% dialyzed FBS (ThermoFisher, 26400044)]. 786-O cells were cultured in RPMI^+pyruvate^ ^+FBS^ [RPMI containing 1 mM pyruvate and 25 mM glucose (ThermoFisher, A1049101) supplemented with 10% non-dialyzed FBS and 1% penicillin/streptomycin] at 37°C in 5% CO_2._ Lenti 786-O cell lines were cultured in RPMI^+pyruvate^ ^+FBS^ in the presence of 1 µg/mL puromycin. All experiments (for the exception of lipidomics with lenti 786-O cells) were performed in RPMI^+pyruvate^ ^+dFBS^ without antibiotics [RPMI containing 1 mM pyruvate and 25 mM glucose supplemented with 10% dialyzed FBS]. Lipidomics experiments with lenti 786-O cell lines were performed in RPMI^+dFBS^ (basal RPMI medium (ThermoFisher, 11879020) without pyruvate and glucose supplemented with 25 mM glucose and 10% dialyzed FBS). Caki-1 cells were cultured in McCoy’s 5A^+FBS^ [pyruvate-free McCoy’s 5A containing 16 mM glucose (ThermoFisher, 16600082) supplemented with 10% FBS and 1% penicillin/streptomycin] at 37°C in 5% CO_2._ Lenti Caki-1 cell lines were cultured in McCoy’s 5A^+FBS^ in the presence of 15 μg/mL blasticidin. All experiments with lenti Caki-1 cell lines were performed in McCoy’s 5A^+dFBS^ medium without antibiotics [pyruvate-free McCoy’s 5A medium supplemented with 10% dialyzed FBS]. All cell lines in this study were mycoplasma free.

### DNA constructs

*Homo sapiens* codon-optimized genes encoding di-domain glycerol-3-phosphate dehydrogenases (GPDHs) from *Dunaliella viridis (GenBank: ACD84644.1)*, *Dunaliella salina (GenBank: AAX56341.1)*, *Chlamydomonas reinhardtii (GenBank: XP_042919880.1),* and *Sphaeroforma arctica (GenBank: XP_014155909.1)* with removed chloroplast targeting sequences (see Supplementary Figure S1) and added a C-terminal linker sequence with a FLAG-tag flanked by EcoRI and AgeI restriction sites were custom synthesized and subcloned into pUC57 vectors by GENEWIZ. After digestion of pUC57 vectors with EcoRI and AgeI restriction enzymes, corresponding DNA fragments were ligated into the pLVX-TetOne-Puro vector (Addgene, Plasmid #124797). For protein characterization studies primers containing BamHI and XhoI restriction sites were used to amplify both _57_*Cr*GPDH_705_ and _87_*Cr*GPDH_705_ constructs from a pUC57 vector which contained *H. sapiens* codon-optimized full-length *C. reinhardtii gpdh* gene with a C-terminal FLAG tag (obtained from GENEWIZ). Resulting gene products were ligated into the pET30a vector (EMD Millipore). Both _57_*Cr*GPDH_705_ and _87_*Cr*GPDH_705_ variants contained an N-terminal Hisx6-tag and a C-terminal FLAG-tag when expressed. All nucleotide sequences were verified by Sanger sequencing (Eton Bioscience, San Diego, CA). To clone Luciferase and _87_*Cr*GPDH_705_ constructs into the pFUW-Blast system for constitutive expression, AgeI and BamHI restriction sites were used.

### Generation of Stable Cell Lines

Lentiviruses were produced by transfecting packaging vectors psPAX2 and pMD2.G together with vectors pLVX-Tet-One-Puro-Luciferase, −_100_*Dv*GPDH_701_, −_98_*Ds*GPDH_701_, −_87_*Cr*GPDH_705_, −_55_*Sa*GPDH_663_ or with vectors pFUW-Blast-Luciferase, −_87_*Cr*GPDH_705_ into HEK293T cells, as previously described^15^. Subsequently, HeLa and Caki-1 Tet-One-Puro lenti cell lines were produced by a single infection with a corresponding pLVX-TetOne-Puro lentivirus followed by selection with 1 µg/mL puromycin. 786-O cells were engineered to constitutively express luciferase or _87_*Cr*GPDH_705_ by infecting cells with corresponding pFUW-Blast-Luciferase or _87_*Cr*GPDH_705_ lentiviruses followed by selection with 15 μg ml/mL blasticidin.

### Expression and Purification of Recombinant *Cr*GPDH

BL21 (DE3) *E. coli* cells transformed with the pET30a-Hisx6-_57_*Cr*GPDH_705_-FLAG or pET30a-Hisx6-_87_*Cr*GPDH_705_-FLAG vectors were grown at 37°C in six 2.8-L flasks, each containing 1 L of Luria-Bertani (LB) medium supplemented with 50 μg/mL kanamycin. When absorbance at 600 nm reached 0.4-0.6, the temperature was decreased to 15°C, and cells were grown for an additional 2 hours before protein expression was induced with 0.1 mM isopropyl β-D-1-thiogalactopyranoside (IPTG). Bacterial cells were subsequently harvested the next morning. All chromatographic steps were performed using an NGC Quest 10 Plus chromatography system (Bio-Rad). For protein purification, the cell pellet was resuspended in 120 mL of the lysis buffer (50 mM Na_2_HPO_4_, pH 8.0, 500 mM NaCl and 30 mM imidazole), and affinity chromatography was performed using a 35 mL Omnifit glass column packed with Ni Sepharose 6 Fast Flow (Cytiva). The purest fractions were pooled and subjected to size-exclusion chromatography on a Superdex 200 Increase 10/300 GL column equilibrated with 50 mM HEPES-NaOH, pH 7.5 and 150 mM NaCl (buffer A). Apparent molecular weights were determined by analytical size-exclusion chromatography on a Superdex 200 Increase 10/300 GL column equilibrated with buffer A by injecting 50 µL of the protein sample (_57_*Cr*GPDH_705_, _87_*Cr*GPDH_705_ or molecular weight standards). A calibration curve was produced using thyroglobulin (669 kDa), ferritin (440 kDa) and beta amylase from sweet potato (200 kDa). The molecular weights of _57_*Cr*GPDH_705_ and _87_*Cr*GPDH_705_ were determined as 419 ± 38 kDa and 301 ± 11 kDa, respectively, indicating that _57_*Cr*GPDH_705_ is a pentamer while _87_*Cr*GPDH_705_ is a tetramer in solution.

### Activity Assays for Recombinant *Cr*GPDH

Activity of the NAD-dependent glycerol-3-phosphate dehydrogenase domain of *Cr*GPDH was monitored in a continuous assay by following the absorbance at 340 nm (ε_340_ =6.2 mM^−1^cm^−1^) using a Cary 3500 UV-Vis spectrophotometer (Agilent Technologies). For the Michaelis–Menten analysis, DHAP was fixed at 5 mM when NADH was varying substrate (2 - 400 µM), and NADH was fixed at 500 µM when DHAP was varying substrate (10 −500 µM). A typical reaction mixture contained in 0.2 mL of buffer A: 5 mM MgCl_2_, NAD(P)H, DHAP and recombinant _87_*Cr*GPDH_705_ (0.29 µg). No enzymatic activity was detected with up to 105 µg of recombinant _87_*Cr*GPDH_705_ with 200 µM NADPH, 5 mM DHAP and 5 mM MgCl_2_in 0.2 mL of buffer A. In addition, the activity of phosphatase (GPP) domain of *Cr*GPDH was monitored by a discontinuous assay when inorganic phosphate (Pi) release was monitored by the malachite green (MG) assay. Because of high background with MG reagent (Sigma, MAK307-1KT), only 50 µM DHAP or Gro3P were used with 100 µM NAD(P)H, 5 mM MgCl_2_ and recombinant enzymes (3.5 - 4.7 μg of _57_*Cr*GPDH_705_ or _87_*Cr*GPDH_705_) in 0.2 mL of buffer A. Typically, an 80 µL aliquot from a reaction mixture was incubated with 20 µL of the MG reagent in a clear 96-well microplate for 30 min, and absorbance was read at 620 nm using BioTek Cytation 10 (Agilent Technologies). In parallel, a calibration curve was produced using known Pi concentrations (0-200 µM).

### Modeling Structures of di-domain GPDHs using AlphaFold

We used web-based AlphaFold interface to model full-length structures of di-domain glycerol-3-phosphate dehydrogenases (GPDHs) from *Dunaliella viridis (GenBank: ACD84644.1)*, *Dunaliella salina (GenBank: AAX56341.1)*, *Chlamydomonas reinhardtii (GenBank: XP_042919880.1),* and *Sphaeroforma arctica (GenBank: XP_014155909.1)* (**Supplementary Figure S2**)^55–57^. We subsequently performed structural alignment using published X-ray structure of *D. salina* di-domain GPDH (PDB#: 6IUY) and our models obtained from the AlphaFold (**Supplementary Figure S1**, **Supplementary Figure S2**). We noticed substantial differences in the N-terminal domain between AlphaFold predicted structures of GPDHs and 6IUY (deposited 6IUY^24^ structure contains gaps in the N-terminal domain). Although authors stated that *D. salina* _99_*Ds*GPDH_699_ variant, they used for structural studies, had a functional N-terminal GPP domain^24^ we think that it was truncated (this explains why our _98_*Ds*GPDH_701_ construct when expressed in HeLa cells accumulated high levels of Gro3P and had a profound proliferation defect) (**Figure 1d, h**; **Supplementary Figure S1**, **Supplementary Figure S3a, h, i**).

### GC-MS Assay for Determination of DHAP, Gro3P and Glycerol

A typical reaction mixture contained in 0.2 mL of buffer A: 5 mM MgCl_2_, 1 mM NAD(P)H, 1 mM DHAP and recombinant _57_*Cr*GPDH_705_ or _87_*Cr*GPDH_705_ (3.5 - 4.7 μg). After incubation of the reaction mixture at 37°C for 10 min, a 100 µL aliquot was quenched with 100 µL of 80/20 % methanol/H_2_O solution containing 100 µM L-norvaline. Samples were dried (Speedvac) along with 7 dilutions of standards containing glycerol, Gro3P and DHAP. Samples and standards were derivatized with 30 µl of isobutylhydroxylamine in pyridine for 20 min at 80°C and with 30 µl of N-tert-butyldimethylsilyl-N-methyltrifluoroacetamide (MTBSTFA) for 60 min at 80°C before they were transferred to GC-MS vials. GC-MS analysis was performed using a Thermo Trace 1610-TSQ 9610 GC-MS/MS instrument fitted with a Thermo TG-S5SILMS column (30 m x 0.25 i.d. x 0.25 μm) (Cancer Metabolism Core, SBP Discovery, La Jolla, CA). The GC instrument was programmed with an injection temperature of 300°C and a 1.0 µl splitless injection. The GC instrument oven temperature was initially 140°C for 3 min, rising to 268°C at 6°C/min, and to 310°C at 60°C/min with a hold at the final temperature for 2 min. GC flow rate with helium carrier gas was 60 cm/s. The GC-MS interface temperature was 300°C and (electron impact) ion source temperature was 200°C, with 70 eV ionization voltage. Standards were run in parallel with samples. Metabolites in samples and standards were detected by MS/MS using corresponding retention times, product ion masses, and collision energies. Sample metabolites were quantified using calibration curves made from the standards in Themo Chromeleon software, and further data processing to adjust for the relative quantities of metabolites in the standards and recovery of the internal standard (norvaline) was done in MS Excel.

### Proliferation Assays

Two thousand HeLa Tet-One-Puro cells were seeded in 0.5 mL of DMEM^+FBS^ in black 96-well microplates with a transparent flat bottom. The next day, media was exchanged with 200 µL of DMEM^+dFBS^ supplemented with 300 ng/mL of Dox and other components as indicated (for hypoxia experiments, microplates were transferred to an incubator set to 0.5% O_2_ or 3.5% O_2_). Experiments with lenti 786-O and Caki-1 cell lines were performed in a similar fashion using specific media for each cell (see Cell Culture section above). On days 1 – 4, cells were fixed using 4% paraformaldehyde in PBS for 15 min, washed twice with PBS, and then kept in 200 µl of PBS supplemented with 1 mg/mL Hoechst and imaged as previously described^15^. Images were processed using Gen5 3.13 software (Agilent Technologies).

### Oxygen Consumption

Oxygen consumption rates (OCR) and acidification rates (ECAR) were measured with the Seahorse XFe96 Flux Analyzer. Four-six thousand HeLa Tet-One-Puro cells per well were seeded in Seahorse 96-well microplates in 80 μL of DMEM^+FBS^. The next day, medium was replaced with 200 μL of DMEM^+dFBS^ ± 300 ng/mL of Dox. Twenty-four hours later, medium was replaced with 200 μL of the Seahorse assay medium [pyruvate free DMEM (US Biological, D9802) supplemented with 10% dialyzed FBS and 25 mM HEPES-KOH with pH adjusted to 7.4]. After basal measurements were collected 6 times, piericidin A or antimycin A were injected as indicated in each experiment and 5 additional measurements were performed. After each assay, the Seahorse 96-well plate was extensively washed with PBS, incubated the SYTOX™ Green Nucleic Acid Stain (ThermoFisher, S7020) for 20 minutes and immediately imaged using BioTek Cytation 10 Confocal Imaging Reader (Agilent Technologies).

### Determination of Total Cellular NADH/NAD^+^ and NADPH/NADP^+^ Ratios

Two-four hundred thousand cells were seeded in 6-cm dishes in 2 mL of cell line-specific media (see section Cell Culture). Twenty-four hours later, media was exchanged with 3 mL of DMEM^+dFBS^ (or RPMI^+pyruvate^ ^+dFBS^/ McCoy’s 5A^+dFBS^) and 300 ng/mL of Dox (no Dox was added to 786-O lenti cell lines which express LUC or _87_*Cr*GPDH_705_ constitutively), and cells were then returned to the incubator. After an additional twenty-four hours, media was exchanged with fresh DMEM^+dFBS^/ RPMI^+pyruvate^ ^+dFBS^/McCoy’s 5A^+dFBS^ that was incubated overnight at 5% CO_2_ supplemented with 300 ng/mL of Dox (for HeLa and Caki-1 lenti lines). Three hours later, 6 cm dishes were placed on ice, washed with 3 mL of ice-cold PBS, and lysed with 0.6 mL of 1:1 mixture of PBS and 1% dodecyltrimethylammonium bromide (DTAB) in 0.2 M NaOH. Samples were processed as previously described^15^, transferred to all-white 96-well microplates, and luminescence was measured over 1.5 hours using Cytation 10 (Agilent Technologies). Only the linear portions of time vs luminescence progress curves for both standards and samples were used in analysis.

### Metabolomic Profiling

One-three hundred thousand HeLa Tet-One-Puro cells were seeded in a 6-well plate in 2 mL of DMEM^+FBS^. Twenty-four hours after seeding, media was exchanged to 2 mL of DMEM^+dFBS^ ± 300 ng/ml Dox. After an additional twenty-four hours, cells were removed from the incubator, placed on ice, lysed (without a PBS wash) with 1 mL of ice cold 80% methanol/20% water solution containing 1.5 µM metabolomics amino acid mix (Cambridge Isotope Laboratories, MSK-A2-1.2). Immediately after, 6-well plates were transferred to −80°C and incubated overnight. The 6-well plates were scraped the next day, and all material was transferred to 1.5 mL Eppendorf tubes and dried in a SpeedVac. Metabolomic profiling analysis using LC-MS was performed as previously described^15^.

### Lipidomics

For lipidomics analysis, two-four hundred thousand cells were seeded in a 6-well plate in 2 mL of cell line-specific media (see Cell Culture section). Twenty-four hours after seeding, media was exchanged to 2 mL of DMEM^+dFBS^ (or RPMI^+dFBS^/McCoy’s 5A^+dFBS^) and 300 ng/mL of Dox (except 786-O lenti cell lines). After an additional twenty-four hours, cells were detached by trypsinization, washed in PBS and frozen until lipids were extracted. For the lipidomic analysis, the frozen cell pellets were extracted using a modified version of the Matyash et al. (2008) method^58^. Briefly, an ice-cold methanol solution containing 0.1 mg/mL butylated hydroxytoluene was added to the pellet, followed by 3 minutes of sonication and 5 minutes of shaking at 15°C and 1000 rpm on a Thermomixer. Next, 900 µL of methyl-tert-butyl ether (MTBE) was added to extract the lipids. The mixture was vortexed for 15 minutes at 4°C and 1000 rpm on the Thermomixer. Phase separation was induced by adding 300 µL of ice-cold water, followed by another 3 minutes of sonication, 15 minutes of vortexing at 15°C and 1000 rpm, and then centrifugation at 21,000 g and 4°C for 15 minutes. The upper organic layer (400 µL) was collected, dried using a Genevac EZ-2.4 Elite evaporator, and stored at −80°C. Separate aliquots were reserved for positive and negative ionization modes. On the day of analysis, the dried lipid extract was resuspended in a 3:2:1 mixture of isopropanol, acetonitrile, and water at room temperature, sonicated for 3 minutes, and vortexed for 15 minutes at 15°C and 2000 rpm. The supernatant was transferred to an LC-MS vial, and a pooled QC sample was prepared from the remaining extracts. Lipids were separated using a Thermo Scientific Accucore C30 column (2.1 × 150 mm, 2.6 µm) connected to a Vanquish Horizon UHPLC system and IQ-X tribrid mass spectrometers. The column was maintained at 45°C, with an injection volume of 4 µL and a flow rate of 0.26 mL/min. Mobile phase A (MPA) consisted of 60/40 acetonitrile/water with 10 mM ammonium formate and 0.1% formic acid, while mobile phase B (MPB) was composed of 89.1/9.9/0.99 isopropanol/acetonitrile/water with the same additives. The chromatographic gradient was set as follows: 0 minutes at 30% B, 2.00 minutes at 43% B, 2.1 minutes at 55% B, 12.00 minutes at 65% B, 18.00 minutes at 85% B, 20.00 minutes at 100% B, maintained at 100% B until 25.00 minutes, then returned to 30% B at 25.1 minutes and held until 30.00 minutes. MS1 parameters were as follows: spray voltage: 3500 V for positive ionization and 2500 V for negative ionization modes, sheath gas: 40, auxiliary gas: 10, sweep gas: 1, ion transfer tube temperature: 300°C, vaporizer temperature: 350°C, orbitrap resolution: 120K, scan range(m/z): 250-2000 for pos, 200-2000 for neg, RF lens(%): 60, automatic gain control (AGC) target: 50%, and a maxIT of 100 milliseconds (ms). Quadrupole isolation and Internal calibration using Easy IC were enabled. Lipids were identified by performing the MS2 and MS3 experiments in the orbitrap mass analyzer. A comprehensive data-dependent HCD MS2 experiment with conditional CID MS2 and MS3 data-acquisition strategy was applied for the in-depth characterization of lipid species. Lipid fragmentation was obtained by the first MS1 data acquisition in full scan mass range (200-2000) followed by the data-dependent (dd) MS2 with the normalized stepped HCD collision energy (%) at 25, 30, 35, OT-30K resolution, maxIT-54 ms. If the HCD fragmentation had the fragment ion-184.0733 m/z (phosphocholine head group), then the same ions were subjected either to ddMS2 CID (fixed collision energy-32%, activation time-10 ms & Q-0.25, 30K resolution, 100 % normalized AGC target and maxIT-54 ms) or CID MS3 scans (fixed collision energy-35%, activation Q-0.25) triggered on the top 3 most intense ions that lost neutral fatty acids plus ammonia (only for triacylglycerols). A total of six injections were made to generate fragmentation data using the AcquireX workflow on the pooled QC samples. The Thermo Scientific LipidSearch software (version 5.0) was used to compile the list of identified lipids (precursor tolerance ± 3 ppm, product tolerance ±5.0 ppm, product threshold-1.0). [M+H] adduct was used to identify and quantify HexCer, SM, SPH, MePC, CoQ, AcylCarnitine, LPC, LPE, PC and PE species while [M+NH] was for the TG, DG, cholesteryl ester species. These identified lipid species were quantified using the Compound Discoverer 3.3 and Skyline^59^ software.

### Transcriptomics

Samples were prepared as previously described^15^. Five hundred thousand HeLa Tet-One-Puro lenti cells were seeded in 10 cm dishes in 10 mL of DMEM^+FBS^. Twenty-four hours later, media was exchanged with 10 mL of DMEM^+dFBS^ ± 300 ng/mL Dox. Twenty-four hours after addition of doxycycline, cells were harvested, and cell pellets were snap frozen. Samples were submitted to GENEWIZ for NGS RNA sequencing, and data analysis was performed as previously described^15^. Differential gene expression analysis was performed using the DESeq2 package in R^15^. Volcano plots were generated to display the global differentially expressed genes across the compared groups using ggplot2 packages in R. Gene ontology clusters were formed using comparisons of significantly changed genes (adjusted p value < 0.05, |log_2_ (Fold change) |> 1) in *Cr*GPDH expressing cells to those in luciferase expressing cells, and the enrichment of gene ontology terms and cnetplots were performed in cluster profiler R package.

### Statistical Analysis

For repeated measurements, statistical analysis was performed using a built-in statistical package in GraphPad Prism 9.3.1. Each experiment presented was repeated independently at least three times with similar results. Exact p values are indicated. Each dot in bar graphs represents a biological replicate. All error bars displayed in the figures represent standard deviation (S.D.).

## Data and Code Availability

RNA-seq data presented in this work are available at the Gene Expression Omnibus database under accession number GEO: XXX.

## Acknowledgments

We thank David Scott (SBP Discovery Cancer Metabolism Core, La Jolla, CA) for technical support. This work was supported by grants from the National Institutes of Health (R35GM142495 and R03CA286706 to VC). The Seahorse XFe96 analyzer in the Saez laboratory (TSRI) was supported by 1S10OD16357. The Metabolomics Platform (RRID: SCR_022932) was supported by the University of Chicago Comprehensive Cancer Center Support Grant (P30 CA014599). This is manuscript 1080 from the Scintillon Institute.

## Contributions

XP performed all experiments with assistance from ALZ, SM and AP. SM performed protein purification and enzyme kinetics experiments. SV and JRC performed LC-MS experiments. HS performed lipidomic profiling experiments. XP, VC and ALZ wrote the manuscript with input from all the authors.

## Supplementary Figures

**Supplementary Figure S1:**
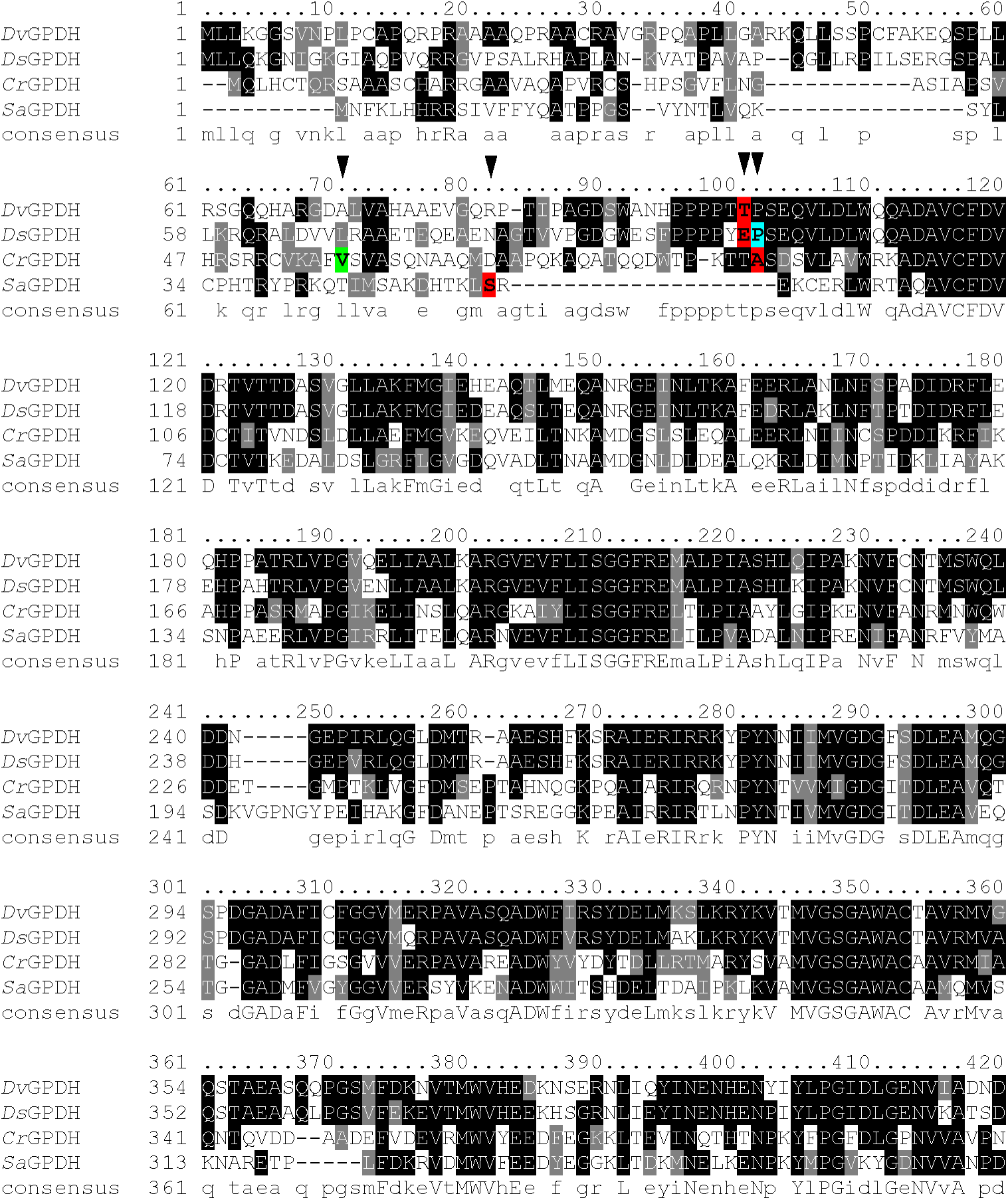
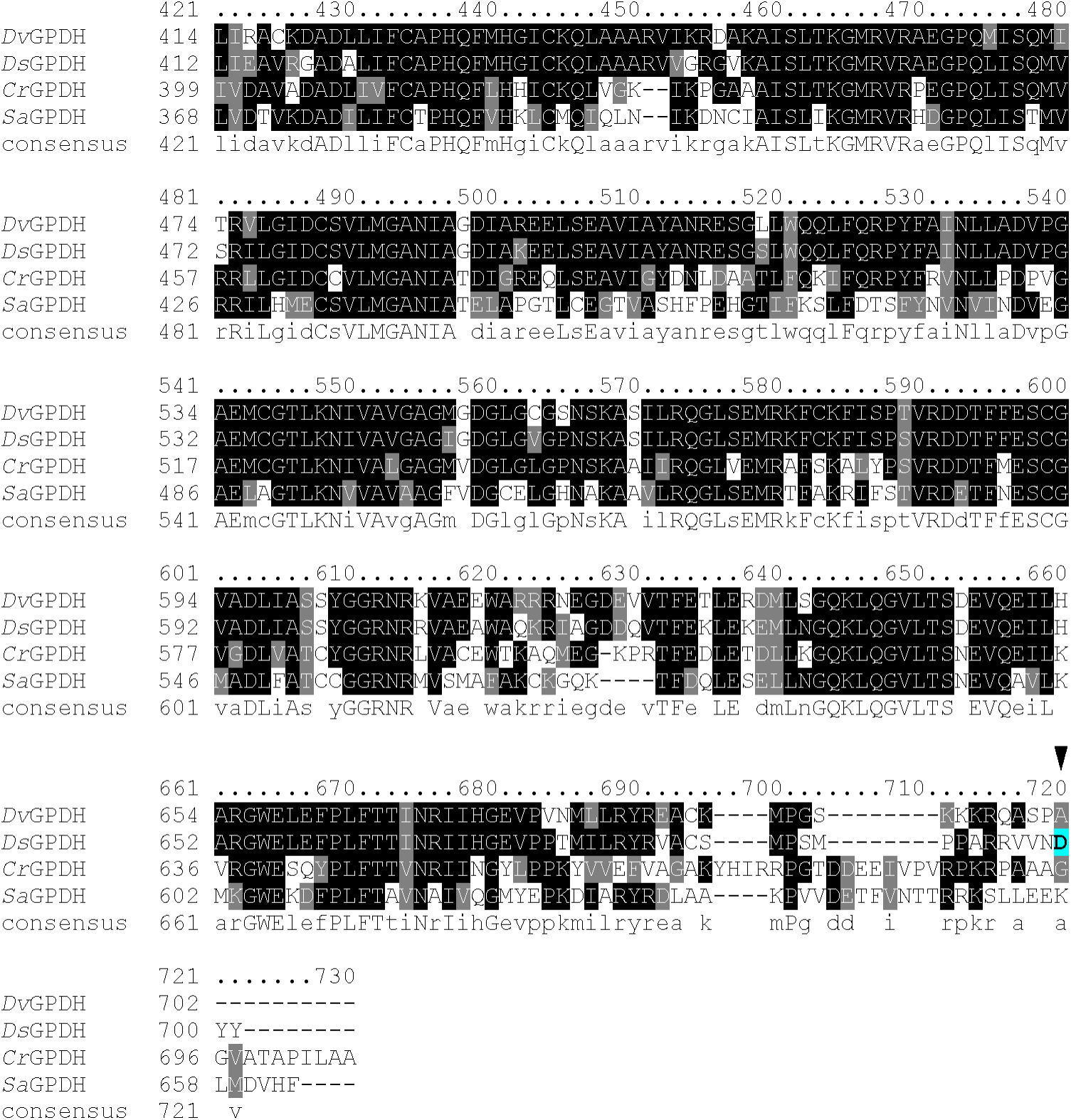
Multiple protein sequence alignment of di-domain glycerol-3-phosphate dehydrogenases (GPDHs) from Dunaliella viridis (GenBank: ACD84644.1), Dunaliella salina (GenBank: AAX56341.1), Chlamydomonas reinhardtii (GenBank: XP_042919880.1) and Sphaeroforma arctica (GenBank: XP_014155909.1). Black inverted triangles and color highlighted amino acids depict: _99_DsGPDH_699_ variant which was previously structurally and biochemically characterized (cyan)^24^; _100_DvGPDH_701_, _98_DsGPDH_701_, _87_CrGPDH_705_, _55_SaGPDH_663_ variants with removed chloroplast targeting sequences (red); and additional _57_CrGPDH_705_ variant (green) which was used to confirm the activity of the N-terminal phosphatase (GPP) domain of CrGPDH.

**Supplementary Figure S2:**
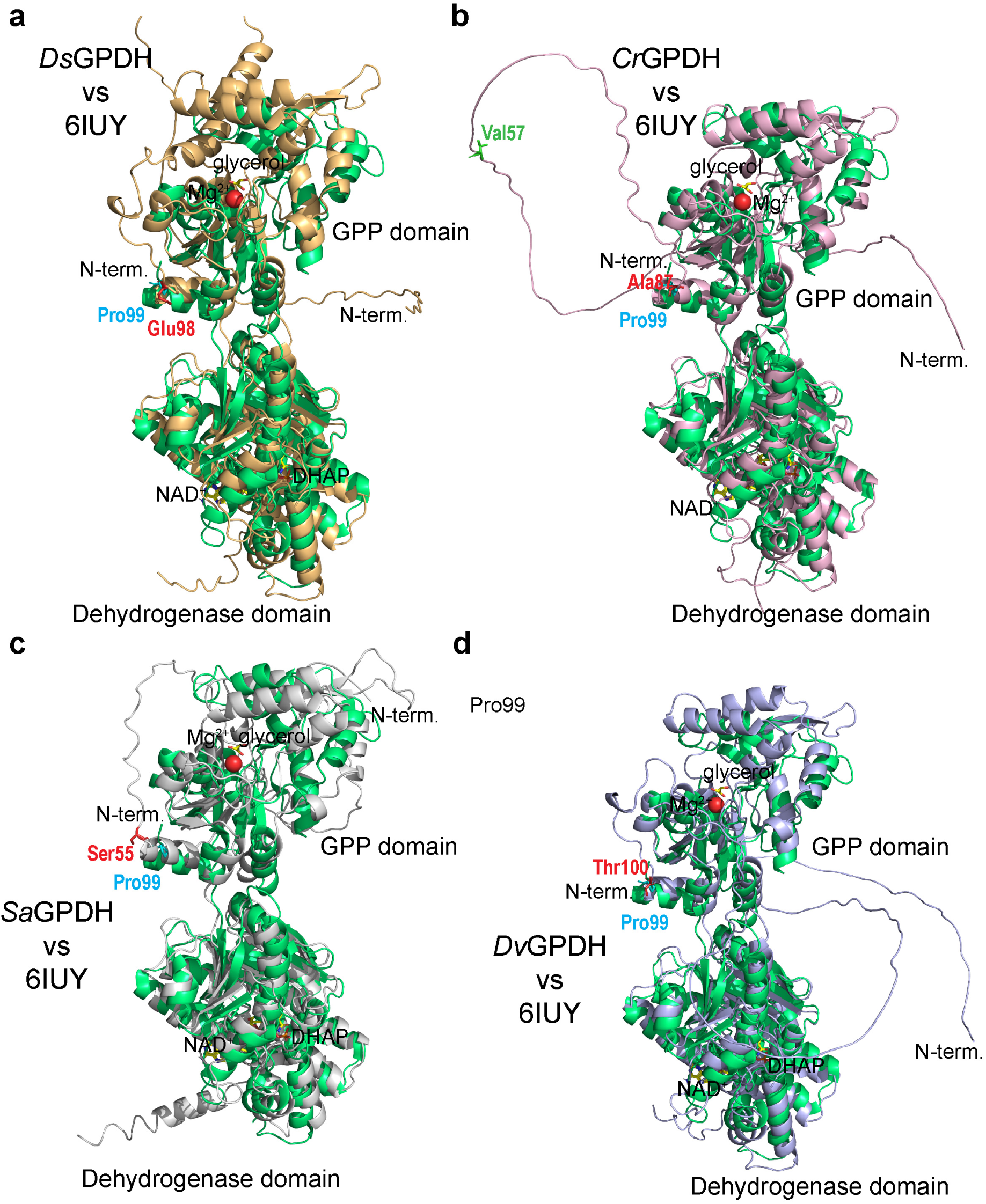
Structural alignment between *D. salina* crystal structure (PDB#: 6IUY) (depicted in green) and models of GPDHs from *D. salina* (**a**)*, C. reinhardtii* (**b**)*, S. arctica* (**c**) and *D. viridis* (**d**) generated by AlphaFold. The X-ray structure (PDB#: 6IUY) in the original study was produced using the _99_*Ds*GPDH_699_ variant (see Supplementary Figure S1) ^24^. NAD^+^, DHAP, glycerol and Mg^2+^ ligands are depicted in 6IUY. Residues representing the N-terminal boundaries are shown in cyan for 6IUY structure; in red for _100_*Dv*GPDH_701_, _98_*Ds*GPDH_701_, _87_*Cr*GPDH_705_, _55_*Sa*GPDH_663_ variants; and in green for _57_*Cr*GPDH_705_ variant (the same color and numbering as in Supplementary Figure S1). We noticed substantial differences in the N-terminal domain between AlphaFold predicted structures of GPDHs and 6IUY (in deposited 6IUY^24^, extensive gaps in the model were present in the N-terminal domain). The major feature of all AlphaFold generated structures is the absence of secondary structure upstream of the GPP domain. This agrees with the presence of extensive chloroplast targeting sequences in all GPDHs at the N-terminus upstream of the GPP domain ^24,60^.

**Supplementary Figure S3:**
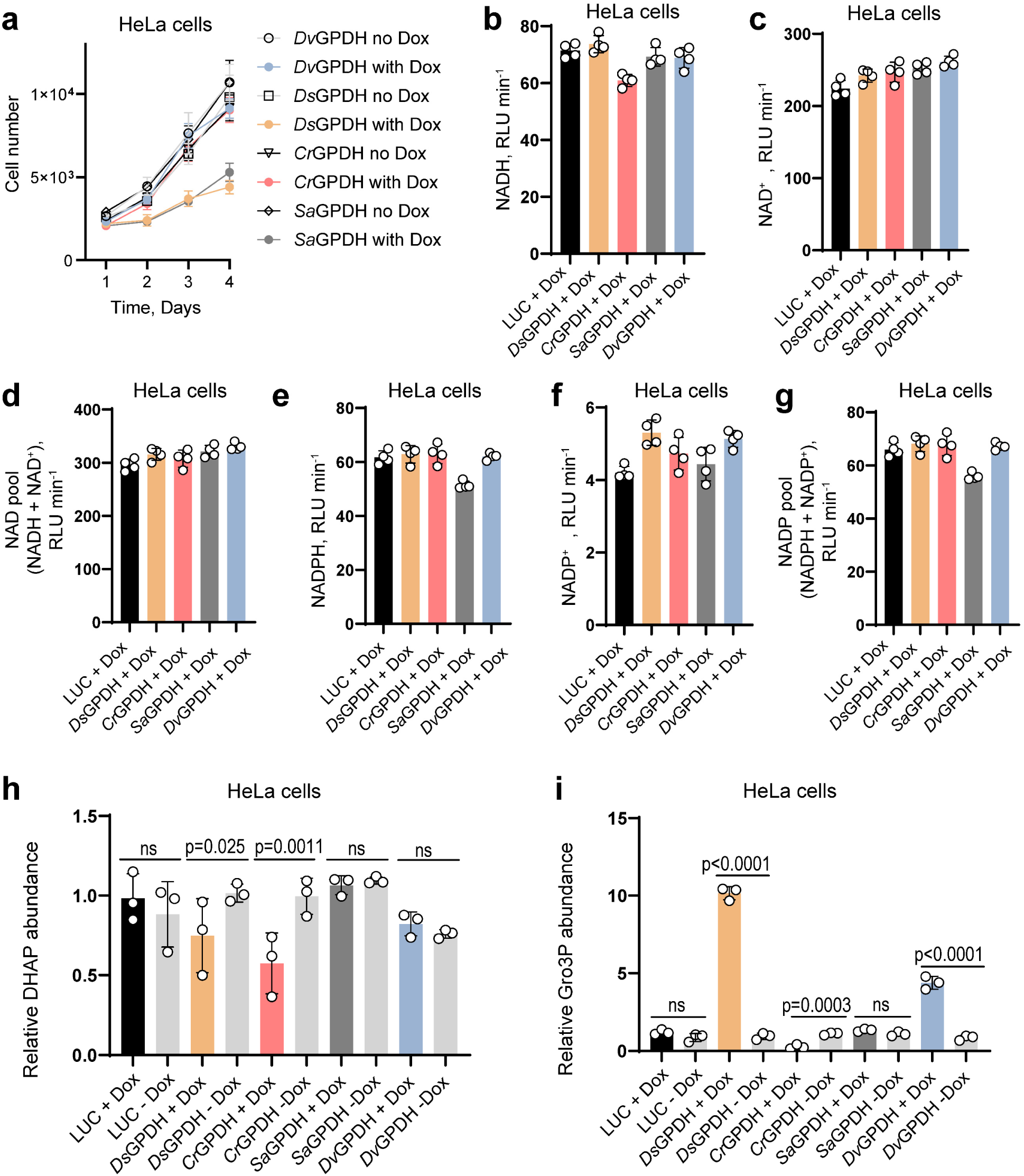
(**a**) Growth curves of HeLa cells infected with pLVX-Tet-One-Puro-GPDHs lentiviruses with and without 300 ng/mL doxycycline (Dox) in pyruvate-free DMEM^+dFBS^. Raw values of luminescence in relative light units (RLU) per minute slopes obtained in enzymatic cycling assays for determination of NADH (**b**), NAD^+^ (**c**), NAD pool (**d**), NADPH (**e**), NADP^+^ (**f**) and NADP pool (**g**) in HeLa cells expressing Luciferase (LUC) and GPDHs from *D. salina, C. reinhardtii, S. arctica* and *D. viridis* under Dox control. Intracellular levels of DHAP (**h**) and Gro3P (**i**) in HeLa cells infected with lentiviruses expressing GPDHs from *D. salina, C. reinhardtii, S. arctica* and *D. viridis* with and without Dox. LUC expressing HeLa cells were used as controls in (b-i). _98_*Ds*GPDH_701_, _87_*Cr*GPDH_705_, _100_*Dv*GPDH_701_ and _55_*Sa*GPDH_663_ variants with removed chloroplast targeting sequences and an added C-terminal FLAG tag were expressed in (a-i). For growth curves in (a), error bars represent mean ± s.d.; n = 6 biologically independent samples. Values are mean ± s.d.; n = 4 in (b-g), n = 3 in (h, j) biologically independent samples. Statistically significant differences in (h, i) were calculated by using a one-way ANOVA followed by uncorrected Fisher′s least significant difference test. NS, no significant difference.

**Supplementary Figure S4:**
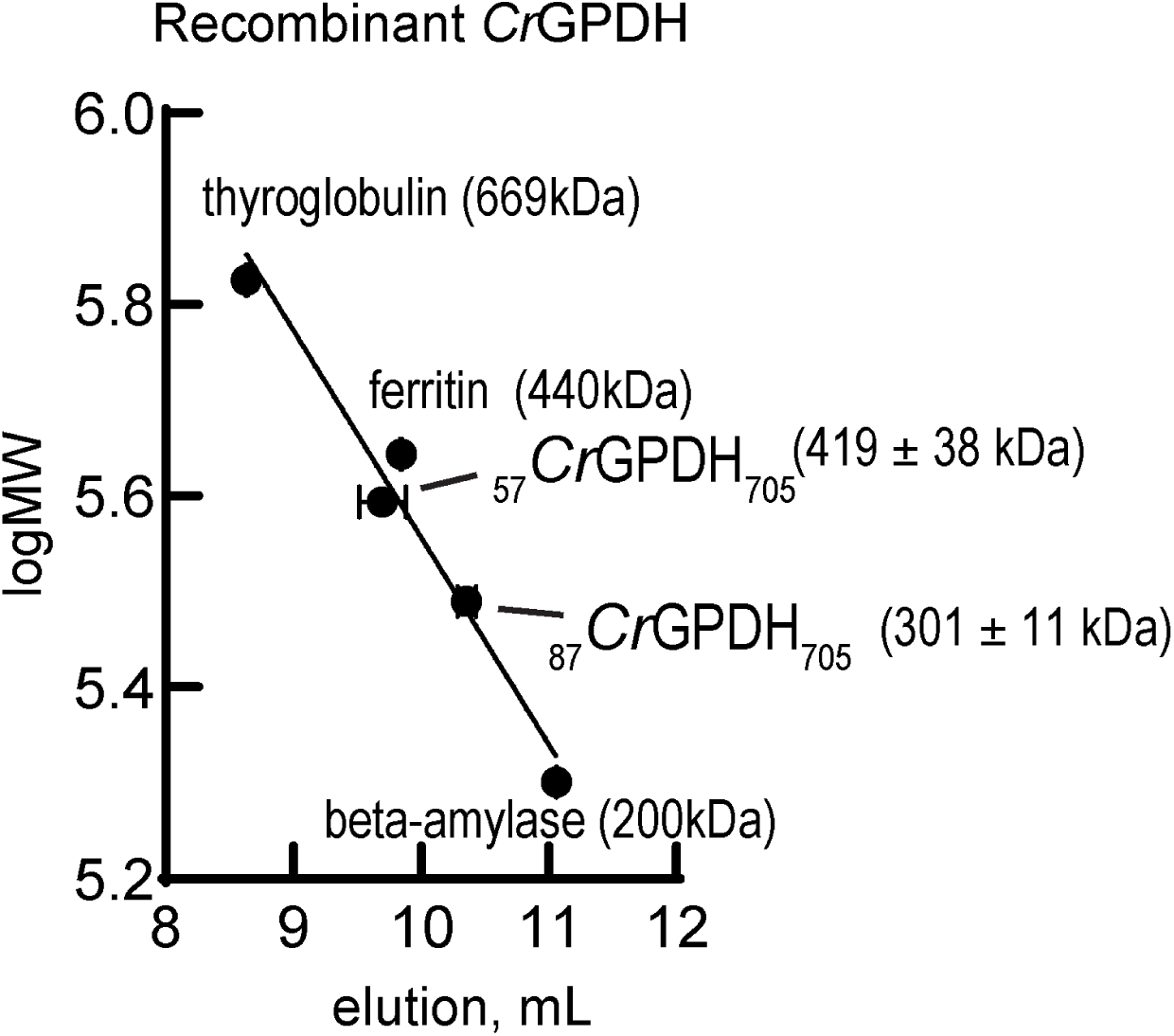
Determination of apparent molecular weight of recombinant _87_*Cr*GPDH_705_ and _57_*Cr*GPDH_705_ variants by size-exclusion chromatography. The calibration curve shown was constructed using thyroglobulin, ferritin and beta amylase, as described under Methods. Calculated apparent molecular weights of _57_*Cr*GPDH_705_ and _87_*Cr*GPDH_705_ were 419 ± 38 kDa and 301 ± 11 kDa, respectively.

**Supplementary Figure S5:**
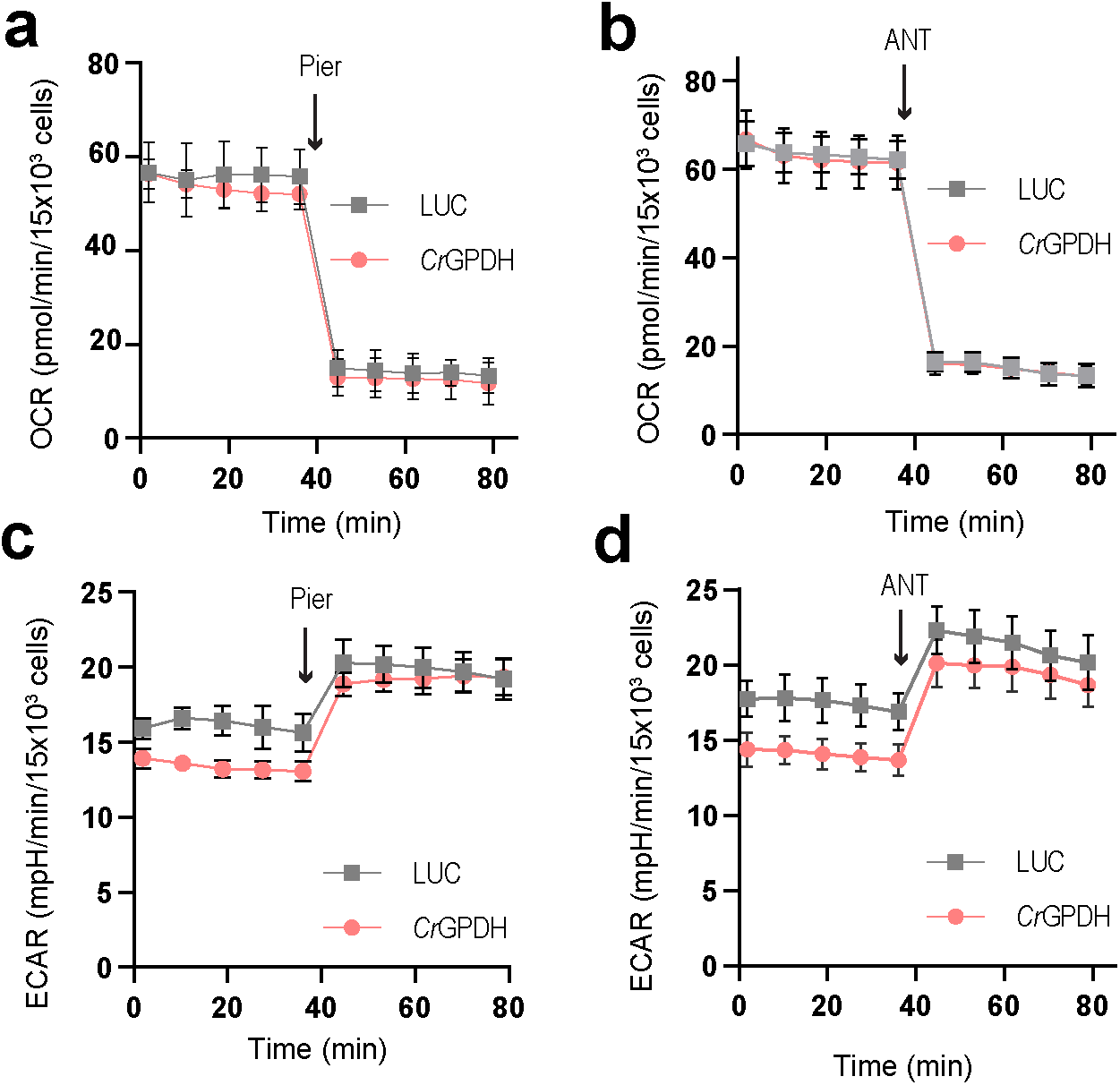
Oxygen consumption rate (OCR) (**a**, **b**) and extracellular acidification rate (ECAR) (**c**, **d**) of HeLa cells expressing LUC and *Cr*GDPH before and after addition of 1 µM piericidin A or 1 µM antimycin A (ANT), measured in pyruvate free HEPES/DMEM^+dFBS^ media. Values are mean ± s.d.; n = 3, 6 in (a), n = 6, 6 in (b), n = 4, 6 in (c), n = 6, 6 in (d) biologically independent samples.

**Supplementary Figure S6:**
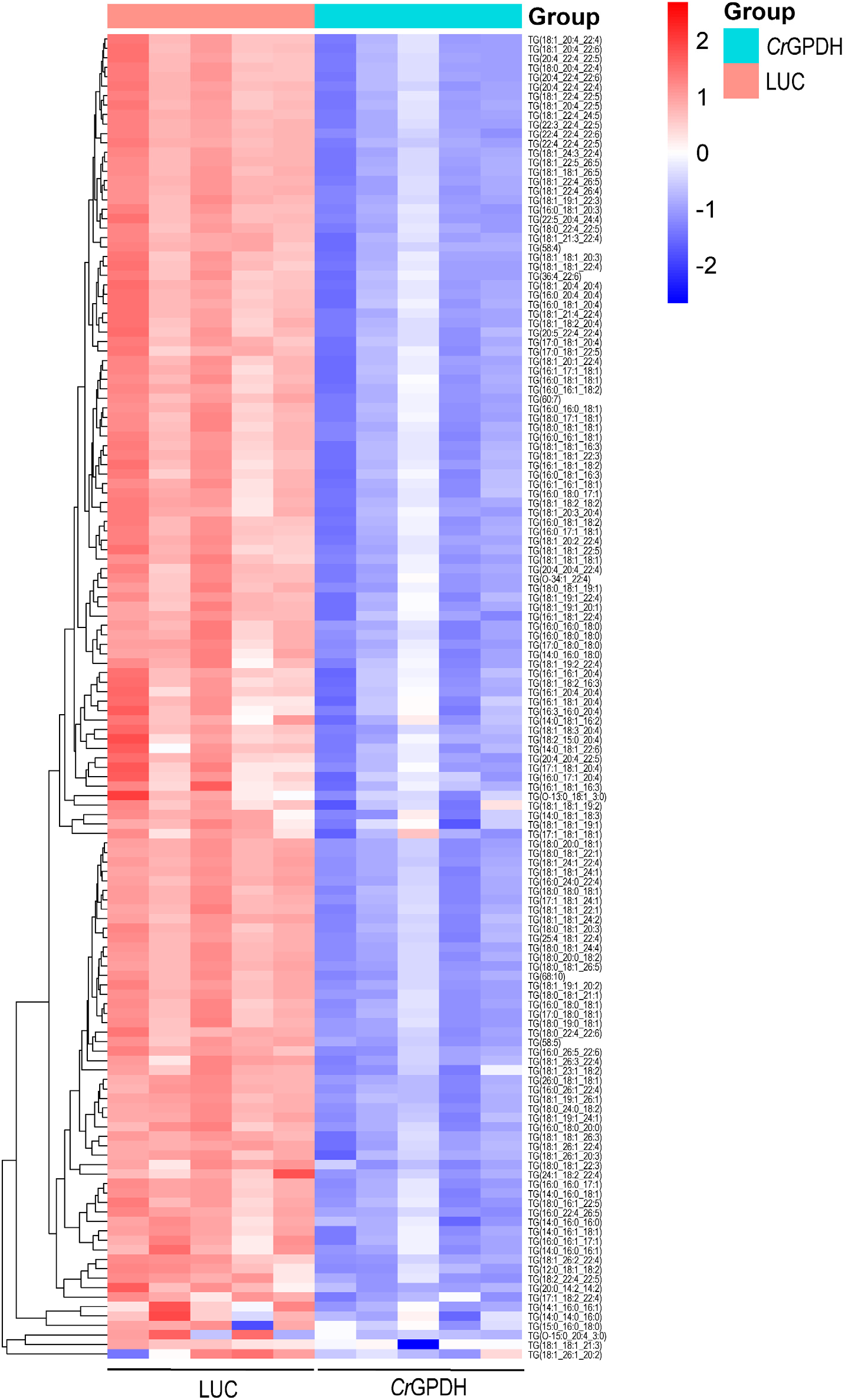
*Cr*GPDH expression in 786-O cells decreases triglycerides levels. In the heatmap, each column represents a biologically independent sample.

**Supplementary Figure S7:**
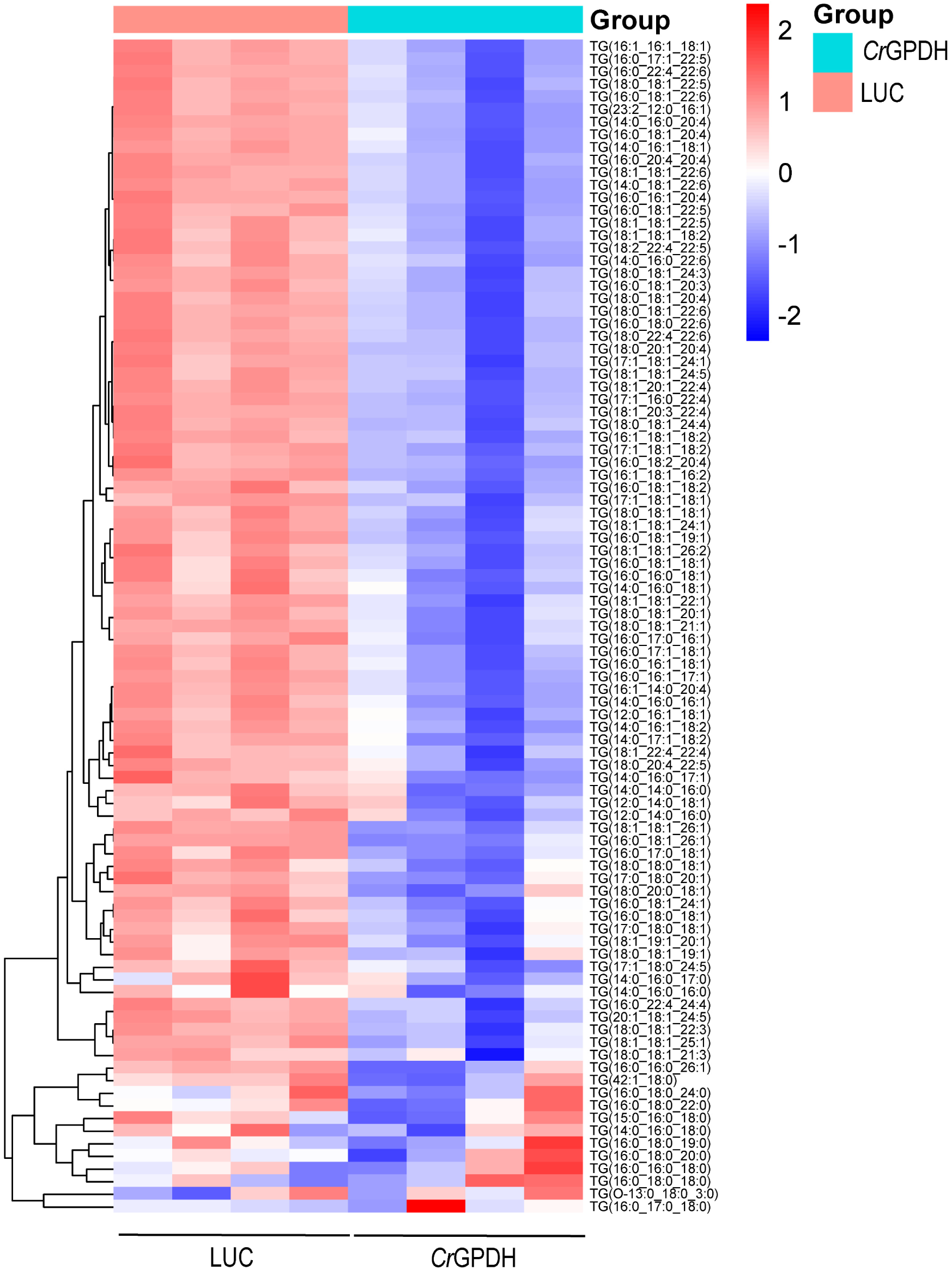
*Cr*GPDH expression in Caki-1 cells decreases triglycerides levels. In the heatmap, each column represents a biologically independent sample.

